# Targeted Photoactivatable Green-Emitting BODIPY Based on Directed Photooxidation Induced Activation and its Application to Live Dynamic Super-Resolution Microscopy

**DOI:** 10.1101/2024.06.20.599858

**Authors:** Lazare Saladin, Valentine Le Berruyer, Maxence Bonnevial, Pascal Didier, Mayeul Collot

## Abstract

Photoactivatable fluorescent probes are valuable tools in bioimaging for tracking cells down to single molecules and for single molecule localization microscopy. For the latter application, green emitting dyes are in demand. We herein developed an efficient green-emitting photoactivatable furanyl-BODIPY (PFB) and we established a new mechanism of photoactivation called Directed Photooxidation Induced Activation (DPIA) where the furan is photo-oxidized in a directed manner by the singlet oxygen produced by the probe. The efficient photoconverter (93-fold fluorescence enhancement at 510 nm, 49% yield conversion) is functionalizable and allowed targeting of several subcellular structures and organelles, which were photoactivated in live cells. Finally, we demonstrated the potential of PFB in super-resolution imaging by performing PhotoActivated Localization Microscopy (PALM) in live cells.

## Introduction

The usefulness of fluorescent probes in bioimaging is no longer to be proven and has considerably contributed to the advancement of knowledge in biology. Among fluorescent probes, those having the ability to irreversibly turn on their fluorescence upon photoirradiation, also called photoactivatable fluorescent probes,^[1], [2], [3]^ mainly found applications in single molecule tracking,^[4], [5]^ and super resolution microscopy based on Photoactivated Localization Microscopy (PALM).^[6], [7], [8], [9]^ In the field, photoactivatable fluorescent proteins are the most used in the bioimaging community, as they can be fused to proteins of interest.^[1], [10]^ However, photoactivatable fluorescent proteins lack versatility and suffer from several limitations,^[11]^ including: 1) Their ability to label only proteins 2) their generally high quantum yields of photoactivation that makes them hard to handle, 3) their limited brightness, and 4) the difficulty to modify their photophysical properties.

In the light of these disadvantages, small molecular probes have emerged as appealing complementary alternatives. Consequently, several photoactivation approaches have been proposed, including: photoelimination of nitrogen (N_2_),^[3],, [12]^ photoinduced disconnection of a spiro-oxazine,^[8]^ photo-uncaging of a fluorescence quencher,^[13], [14], [15], [16]^ nitrosamine caging groups,^[17]^ diazoketone-caging,^[18]^ light-induced protonation of rhodamine,^[4]^ as well as radical reaction on xanthone dyes.^[19], [20]^ Interestingly, photoactivatable dyes based on photooxidation mediated by ^1^O_2_ led to general and efficient approaches to develop visible-light-activated probes.^[5], [21], [22]^ In the field of single molecule localization microscopy (SMLM), fluorescent probes emitting in the green region of the visible spectrum remain rare. Hence, the latter are in high demand as they would allow multicolor super resolution microscopy in combination with existing red-shifted probes.

BODIPYs are versatile green emitting dyes,^[23]^ and proved their efficiency in bioimaging owing to their brightness and relative photostability.^[24]^ Consequently, several green-emitting photoactivatable dyes based on BODIPYs have been described in the literature (Figure 1). Although some examples report impressive fluorescence enhancements upon photoconversion (up to 1250-fold),^[16], [25]^ these require UV irradiation which is detrimental for live cells experiments. However, BODIPYs photoactivated by visible light also have been reported. In 2018, Wijesooriya *et al.* used the photocage properties of alkyl boron dipyrromethene to develop a photoactivatable BODIPY with a fluorescent quantum yield enhancement of 6-fold.^[26]^ Recently, Zhang *et al.* reported a series of visible light-photoactivatable probes based on PET quenching.^[5]^ Among their examples a photoactivatable BODIPY was developed with a fluorescence enhancement of 37-fold without any production of side products compared to examples mentioned above.^[5]^ Recently we reported a mechanism called Directed Photooxidation Induced Conversion (DPIC) allowing to obtain dual-color emissive photoconverters.^[27], [28], [29]^ This mechanism is based on the coupling of an Aromatic Singlet Oxygen Reactive Moiety (ASORM) to a fluorophore. Under irradiation, the obtained red-shifted dye produces singlet oxygen (^1^O_2_), which photo-oxidizes the ASORM in a directed manner. This process disrupts the conjugation of the dye with the ASORM, leading to a hypsochromic shift in both absorption and emission.^[27]^ This approach, using furan or pyrrole as ASORMs, was successfully applied to coumarins,^[27], [28]^ and BODIPYs (Figure 2A). ^[29]^Indeed, when a pyrrole moiety was added in α position of BODIPYs (Figure 2A), both pyrrolyl-BODIPYs and their converted forms possessed high brightness along with a large shift upon conversion (>70 nm). Noteworthy, this mechanism only requires visible light and does not generate side products upon photoconversion. Herein we hypothesized that placing an ASORM in β position of BODIPY will 1) provide weakly emissive dye as reported in the literature where an aromatic group is added at this position,^[30], [31]^ and 2) allow the dearomatization of the ASORM upon directed photooxidation, thus leading to the photoactivation of the BODIPY(Figure 2).^[27], [28], [29]^ In this work, we showed how this approach proved successful, leading to an efficient photoactivatable green emitting probe that can be targeted to various cellular localizations to perform selective photoactivation as well as dynamic super resolution imaging based on Single Molecule Localization Microscopy (SMLM).

**Figure 1.**
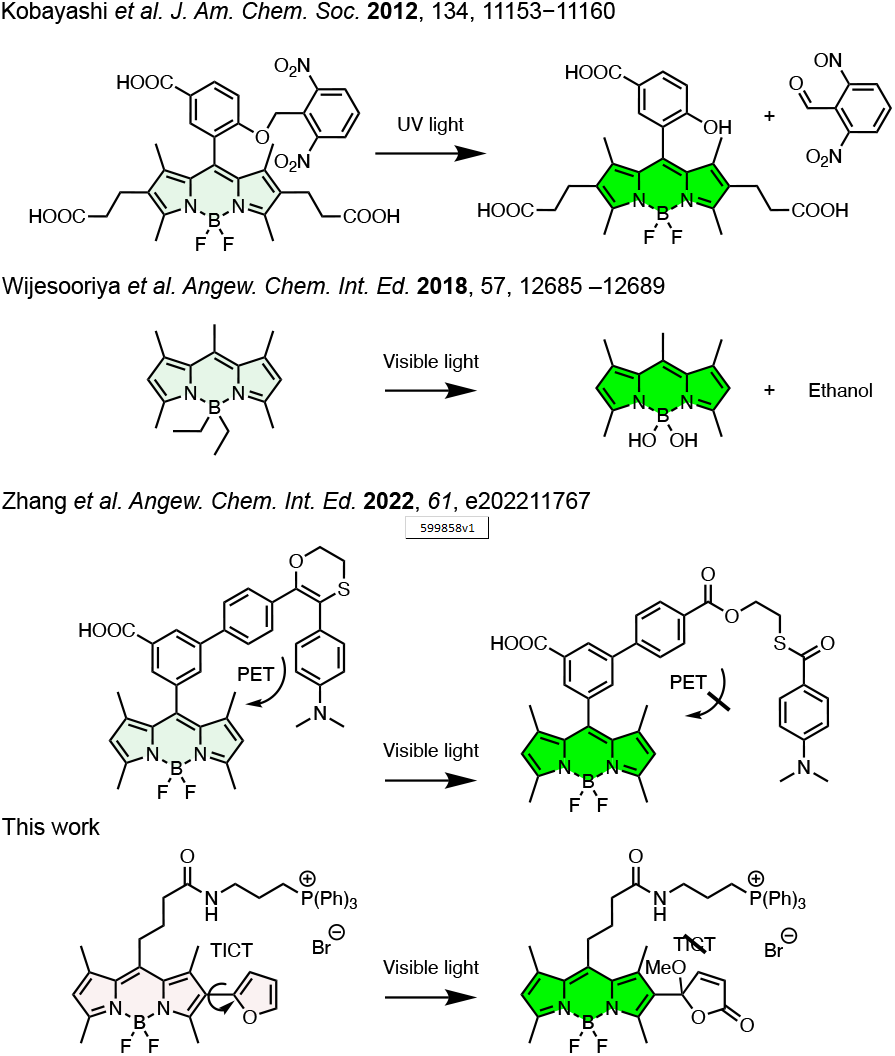
Examples of photoactivatable green emitting BODIPYs given in the literature and based on different mechanisms and the presented work based on Directed Photooxidation Induced Activation.

**Figure 2.**
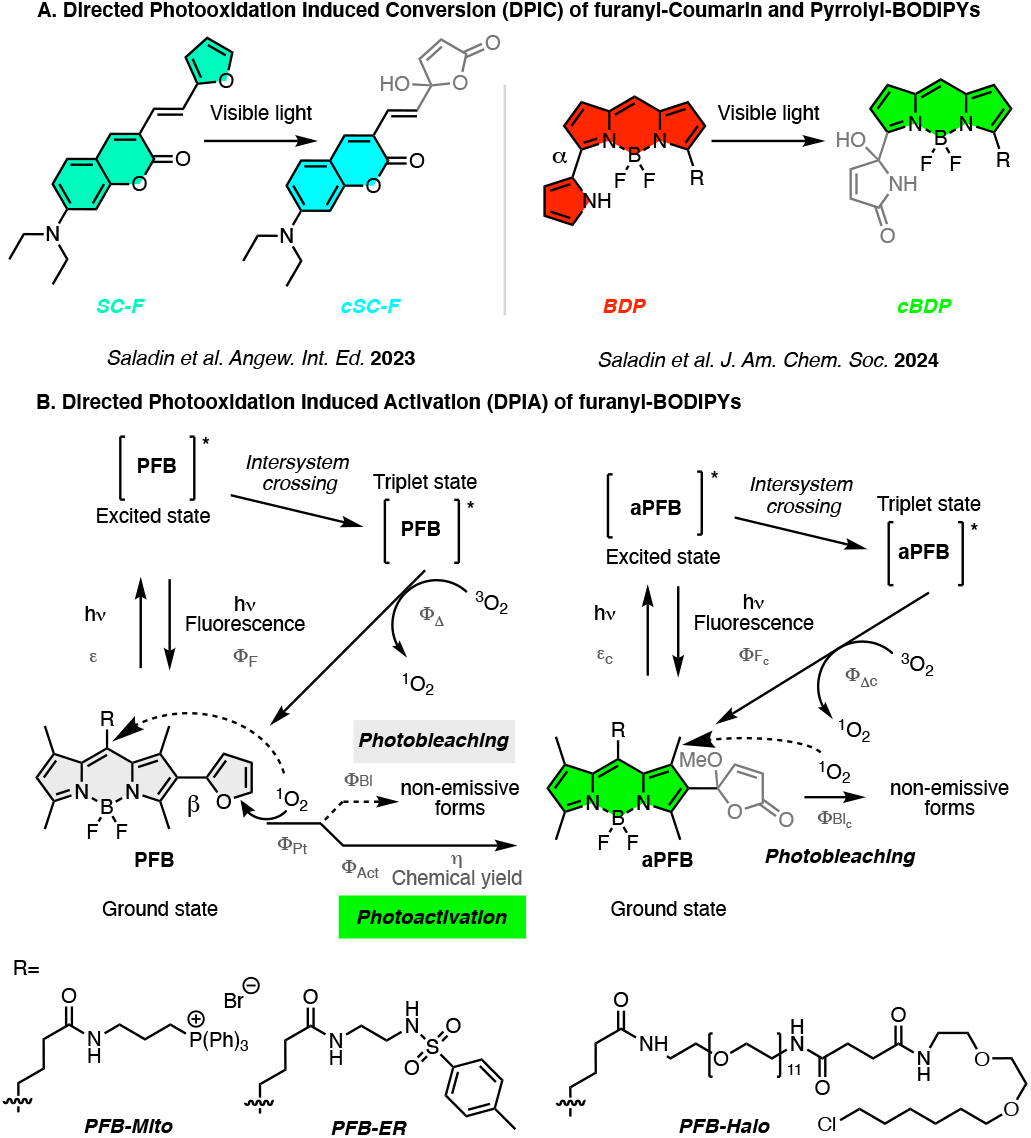
(A) Previous works on photoconvertible probes obtained through Directed Photooxidation Induced Conversion (DPIC) using furan and pyrrole as Aromatic Singlet Oxygen Reactive Moiety (ASORM). (B) Proposed mechanism of photoactivation of the Photoactivatable Furanyl-BODIPY (PFB) through Directed Photooxidation Induced Activation (DPIA). Most of constants (in grey) were determined (Table 1) and supported the proposed mechanism. Various R groups were used to target PFB and led to three different probes.

## Results and Discussion

### Design and synthesis of PFB

As discussed above, we assumed that a photoactivatable green-emitting fluorescent probe could be obtained from the coupling of an ASORM in β position of a BODIPY (Figure 2). Moreover, to extend its applications, we designed a functionalizable probe. Consequently, the tetramethyl BODIPY 1, possessing an ester ended aliphatic chains in *meso* position was chosen as our scaffold (Scheme 1). Furan was chosen as the aromatic singlet oxygen reactive moiety (ASORM). This choice was driven by the better chemical yield of conversion (97 %) provided by furan compared to pyrrole (30 %) when applied to coumarin dyes.^[27]^ Compound 1 was thus iodinated using chloroiodide providing 2, and the furan was introduced in β position through a Suzuki coupling to obtain 3. To make it functionalizable, 3 was saponified to give access to 4, which was coupled to the amine 5 to obtain PFB-Mito, bearing a triphenylphosphonium group known as an efficient mitochondrial targeting moiety.^[32]^ In parallel, 1 was saponified giving rise to acid 6 which was coupled to the mitochondrial targeting moiety 5 to obtain a model BODIPY (MB-Mito). The latter served to evaluate the effect of the furan moiety in the photoactivation process as well as to have a model of photoproduct after photoactivation (See SI).

**Scheme 1.**
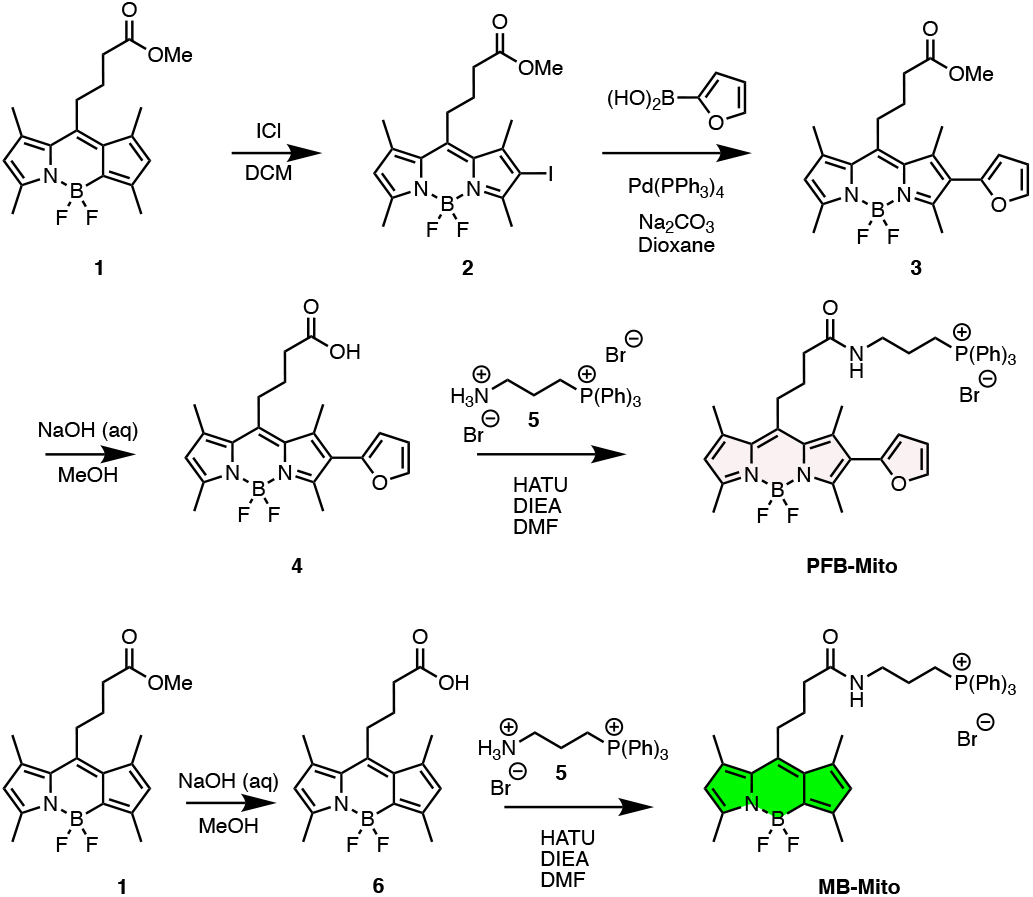
Synthesis of PFB-Mito and the model of photoproduct MB-Mito.

**Table 1.** Photophysical properties of PFB-Mito, its emissive activated photoproduct aPFB-Mito and the model of photoactivated form MB-Mito. 1 µM in methanol.

| Dye | $\lambda_{\text{Abs}}$<br>(nm) | $\epsilon$<br>($\text{M}^{-1}\cdot\text{cm}^{-1}$) | $\lambda_{\text{Em}}$<br>(nm) | $\Phi_{\text{fl}}$<br>(%)<sup>a</sup> | $\tau$<br>(ns)<sup>b</sup> | $\Phi_{\text{Pt}}$ (%)<sup>c</sup> | $\Phi_{\text{Act}}$ (%)<sup>d</sup> | $\Phi_{\text{Bl}}$ (%)<sup>e</sup> | $\eta$ (%)<sup>f</sup> | $\Phi_{\Delta}$<br>(%)<sup>g</sup> |
| --- | --- | --- | --- | --- | --- | --- | --- | --- | --- | --- |
| PFB-Mito | 508 | 40,000 | 628 | 3 | $0.3740 \pm 0.0008$ | $3.5 \pm 0.6 \cdot 10^{-2}$ | $1.7 \pm 0.3 \cdot 10^{-3}$ | $1.8 \pm 0.4 \cdot 10^{-3}$ | $49 \pm 4$ | 4 |
| aPFB-Mito | 501 | N/A<sup>h</sup> | 504 | 34<sup>i</sup> | $3.53 \pm 0.02$ | $1.4 \pm 0.3 \cdot 10^{-4}$ | N/A<sup>j</sup> | $1.4 \pm 0.3 \cdot 10^{-4}$ | N/A<sup>j</sup> | / |
| MB-Mito | 497 | 62,300 | 513 | 100 | $5.50 \pm 0.01$ | $2.0 \pm 0.1 \cdot 10^{-3}$ | N/A<sup>j</sup> | $2.0 \pm 0.1 \cdot 10^{-3}$ | N/A<sup>j</sup> | / |

<sup>a</sup> Fluorescence quantum yield<sup>b</sup> Fluorescence lifetime<sup>c</sup> Quantum yield of phototransformation<sup>d</sup> Quantum yield of photoactivation<sup>e</sup> Quantum yield of photobleaching<sup>f</sup> Chemical yield of photoactivation<sup>g</sup> Quantum yield of $^1\text{O}_2$ generation<sup>h</sup> Non-isolated photoproduct<sup>i</sup> Determined after the photoactivation of PFB-Mito (5 μM), on non-isolated photoproducts<sup>j</sup> Does not photoactivate

### Photophysical properties

Once synthesized and characterized (see SI), the photophysical properties of PFB-Mito were studied. First, impact of the furan moiety when placed in β-position of the BODIPY was observed when the absorption and emission spectra of PFB-Mito and MB-Mito were compared (Figure 3A). PFB-Mito displayed a broad, less intense, and red-shifted emission spectrum (λ_Em max_ = 628 nm) compared to MB-Mito suggesting multiple vibrational transitions probably owing to the dissymmetric and twisted structure induced by the furan moiety. Interestingly, PFB-Mito conserved a high excitation at the common 488 nm laser line (55%, figure 3A), which will allow an efficient photoactivation during imaging. Whereas the model MB-Mito possesses a quantitative fluorescence quantum yield of 1, PFB-Mito displayed an impressive decrease of brightness owing to its lowered extinction coefficient (40,000 M^-1^ cm^-1^) and its extinguished fluorescence quantum yield of 0.03 (Table 1). To determine the mechanism underlying the quenching of PFB-Mito, the solvatochromic behavior of PFB-Mito was assessed (Figure 3B) and suggested that its lowered fluorescent quantum yield occurred through the twisted intramolecular charge transfer (TICT) mechanism.^[33], [34], [35]^ This hypothesis was further confirmed by time-resolved fluorescence decay measurements for which PFB-Mito displayed a decreased fluorescence lifetime of 0.37 ns compared to MB-Mito, 5.50 ns (Table 1, figure S1-2), in agreement with their respective fluorescence quantum yields (Table 1).

**Figure 3.**
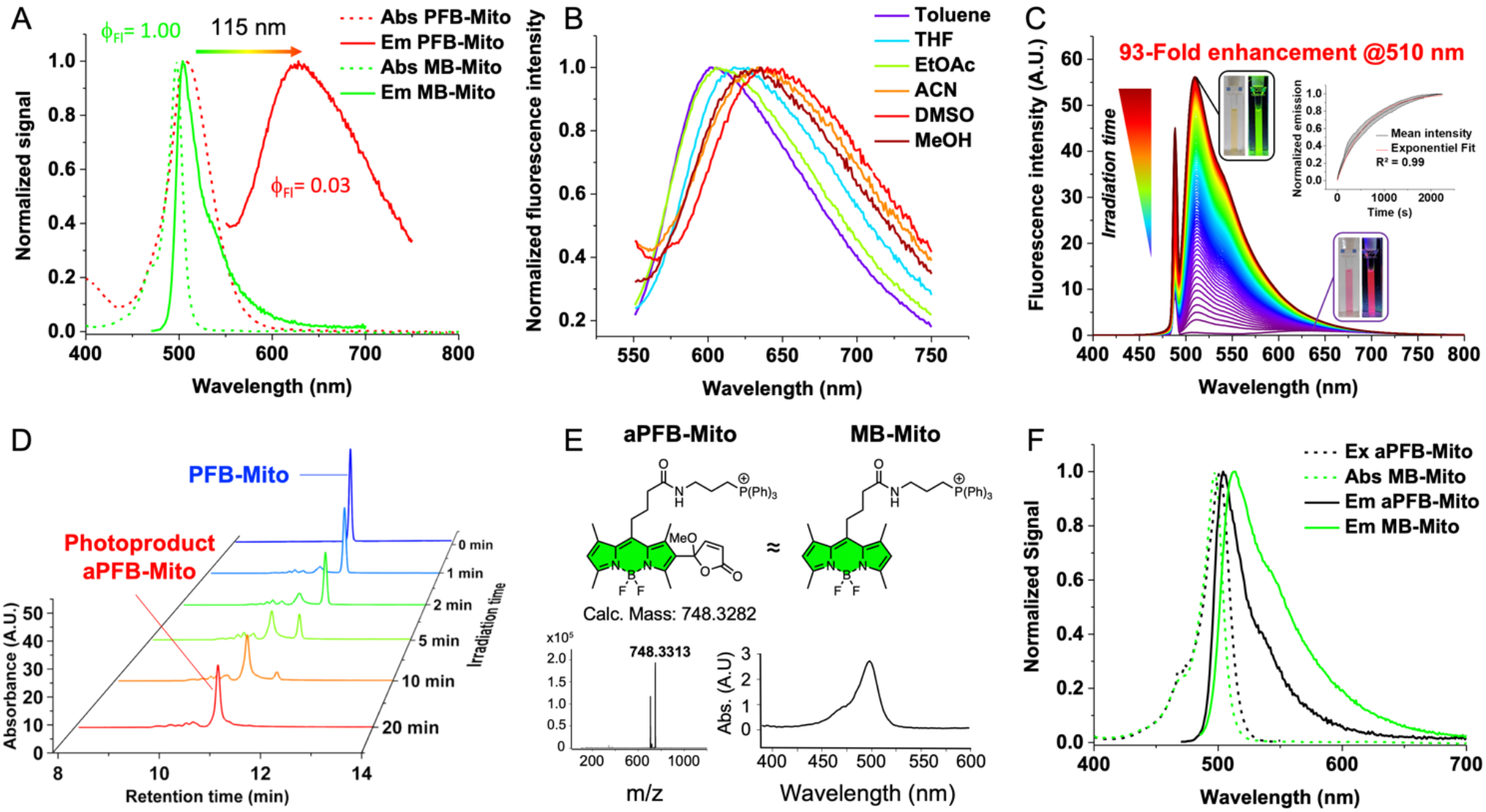
Photophysical and photoactivation studies of PFB-Mito and its comparison with MB-Mito. (A) Absorption and emission spectra of PFB-Mito and MB-Mito showing the impact of the furanyl moiety in β position. (B) Normalized emission spectra of PFB-Mito in solvents with increasing polarity depicting the positive solvatochromism of PFB-Mito and suggesting a quenching through TICT. (C) Emission spectra upon photoactivation of PFB-Mito into aPFB-Mito (excitation was at 488 nm, 160 mW.cm^-2^), inset is the plot of fluorescence intensity at 508 nm over time (red curve is the exponential fit) used to determine the photoactivation constants. (D) HPLC traces of PFB-Mito over the time upon irradiation at 488 nm (the absorbance signal was monitored at 502 nm). (E) Characterization (mass and absorption spectrum) of the main photoproduct aPFB-Mito arising from the photooxidation and its similarity with the model MB-Mito. (F) Excitation and emission spectra of aPFB-Mito overlaid with absorption and emission of MB-Mito showing that the emissive photoproduct was similar to the model.

### Photoactivation properties

To assess its photoactivation properties, a methanolic solution of PFB-Mito was continuously irradiated by a 488 nm laser and the fluorescence emission spectra were recorded over time (Figure 3 C). Rapidly the feint emission band at 628 nm was replaced by an intense and sharp band centered at 510 nm with an impressive fluorescence enhancement of 93-fold (Figure 3C). The fluorescence intensity at 510 nm over time was plotted (Inset Figure 3C) and provided a quantum yield of phototransformation of 3.5 ± 0.6 10^-2^ % (see methods section in Supplementary Information). After photoactivation, a fluorescence quantum yield of the activated form, called aPFB-Mito, was estimated at 34% (Table 1), which is likely minimized by the formation of non-emissive absorbing forms. Additionally, the fluorescence lifetime increased from 0.37 ns to 3.53 ns for PFB-Mito and aPFB-Mito respectively (Table 1, figure S3), suggesting a recovery of brightness through the cancellation of the TICT process. The photoactivation process was monitored over time by HPLC and showed that PFB-Mito (retention time: 11.2 min) progressively disappeared upon irradiation whereas a more polar major photoproduct absorbing at 501 nm progressively appeared (retention time: 10.8 min) (Figure 3D-E). The mass analysis of the major photoproduct (M+O+CH_3_O) was in accordance with the photochemistry of furan derivatives,^[36]^ and with our proposed mechanism where the furan was oxidized by singlet oxygen followed by a solvolysis of the methanol giving rise to a furanone, which is no longer conjugated to the BODIPY (Figure 3E, figure S4). Whereas the emission spectrum of the photoactivated form, aPFB-Mito, was slightly red-shifted (Δλ_Em max:_ 9 nm) and broader compared to our model MB-Mito, their sharp excitation spectra superimposed well (Figure 3F). These results reinforced MB-Mito as a suitable model of photoactivated photoproduct and the latter was thus used to determine key photoconversion parameters.

Based on our model and calculations, PFB-Mito possesses a high chemical yield of photoactivation (η) of 49 ± 4 %, in line with our HPLC analysis displaying a minimized number of photoproducts (Figure 3D). Consequently, this chemical yield (η) of photoactivation indicated similar quantum yields of photobleaching and of photoactivation (1.8 ± 0.4 10^-3^ % and 1.7 ± 0.4 10^-3^%, respectively). In plain words: PFB-Mito bleaches as fast as it photoactivates toward aPFB-Mito upon 488 nm irradiation. PFB is thus less photoreactive compared to photomodulable proteins displaying phototransformation quantum yield ranging from 10^-1^ to 10^-2^ %,^[37], [38]^ and would thus avoid undesired triggering of photoactivation during sample manipulations in microscopy experiments.

To decipher which reactive oxygen species (ROS) could trigger the photooxidation of PFB, its reactivity toward various ROS has been assessed in spectroscopy. The results indicated that PFB, in addition to light, could be activated at its ground state by singlet oxygen (Figure S5), which is in line with our previous reports,^[27],[28]^ as well as with studies related to the reactivity of furan.^[36]^ Since PFB can get activated under irradiation of light, its quantum yield of singlet oxygen generation (β) was determined and gave a value of 4%, which is slightly higher than our photoconvertible BODIPYs based on DPIC.^[29]^ Hence, these experiments confirmed our proposed mechanism of Directed Photooxidation Induced Activation (DPIA) where PFB generates the singlet oxygen necessary to react with the furan moiety responsible for the photoactivation.

Finally, the photostability of the photoactivated form aPFB-Mito was assessed as it is a key parameter for bioimaging tracking or SMLM applications. Surprisingly aPFB-Mito displayed a high photostability depicted by a 14-fold lower quantum yield of photobleaching (β_Bl_ = 1.4 10^-4^ %) compared to the model MB-Mito (β_Bl_ = 2.0 10^-3^ %).

Overall PFB displayed both appealing photophysical and photoactivation properties including: a high compatibility with visible light excitation using the widely used 488 nm laser line, a balanced photoconversion rate combined with a high yield of conversion, as well as bright and photostable photoactivated photoproduct.

### Cellular Imaging

To evaluate the efficiency of PFB as a photoactivatable probe in bioimaging, PFB-Mito was first incubated with HeLa cells at 200 nM. As expected and given the low brightness of PFB-Mito, no signal was observed in the green channel (Figure 4A). Cells contours were localized by a plasma membrane staining using MemBright 640 and after zooming on a single cell followed by consecutive scans with the 488 nm laser line, the cytoplasmic intensity of fluorescence increased to finally reach a plateau after several scans (≈20) (Figure 4B, movie 1). The experiment has been repeated on several cells of the same image field without triggering the photoactivation of surrounding cells thus proving an excellent spatiotemporal control of the photoactivation (Figure 4A-B, S6).

**Figure 4.**
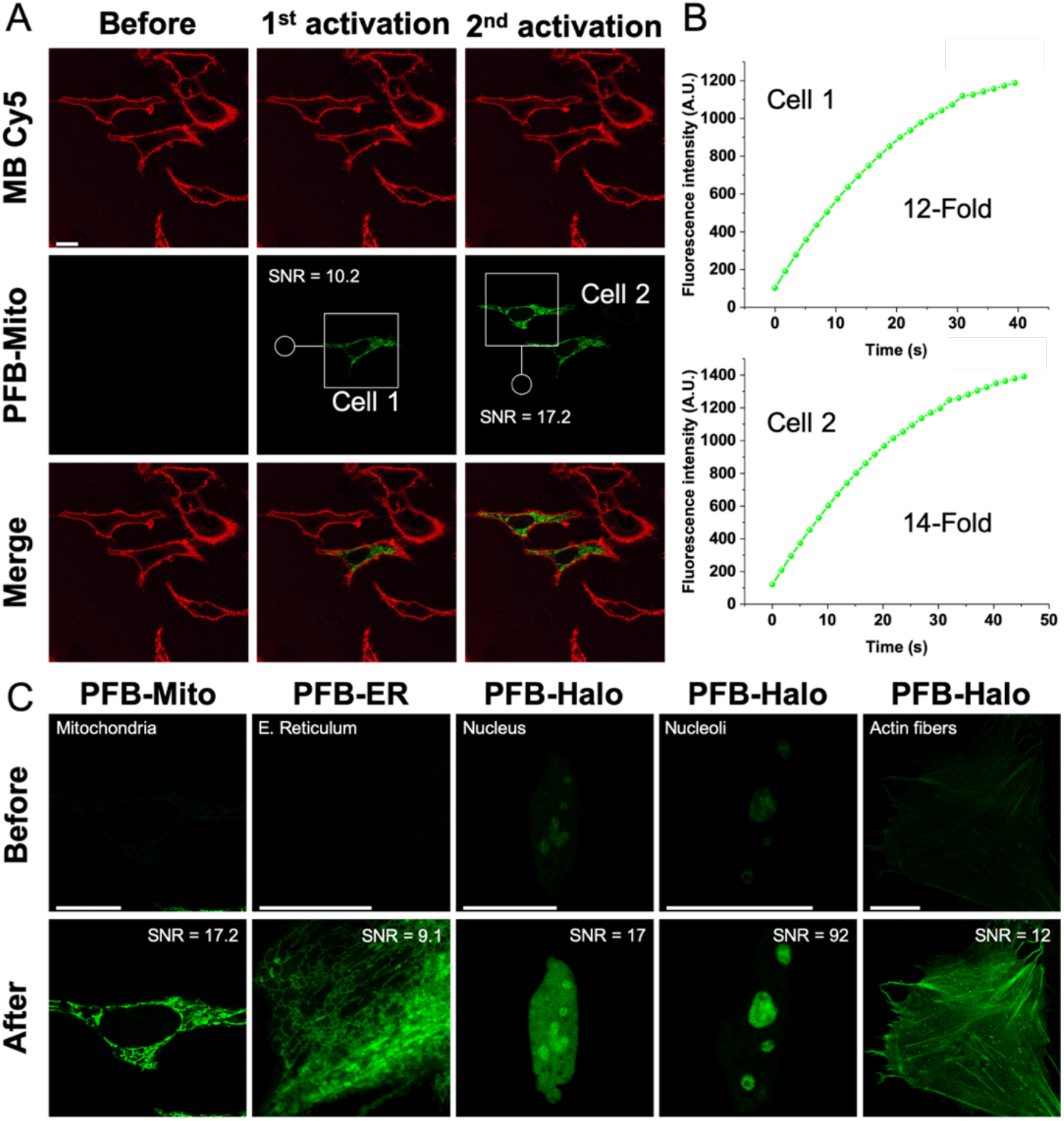
Photoactivation of PFB targeted to various organelles. (A) Laser scanning confocal microscopy images of HeLa cells incubated with PFB-Mito (200 nM) and MemBright 640 (200 nM). Single cells were sequentially and selectively photoactivated upon scanning on the region of interest. (B) Increase of the mean fluorescence intensity upon scanning at 488 nm for individual cells. (C) Laser scanning microscopy images of HeLa cells stained with PFB-Mito, PFB-ER and PFB-Halo before (top) and after photoactivation (bottom) showing the efficient activation in various organelles. SNR is the measured Signal-to-Noise Ratio. Scale bar is 20 µm.

The selective staining of mitochondria in cells by aPFB-Mito was confirmed using Mito-Tracker™ deep red as a co-staining marker with a Pearson’s coefficient of 0.77 (Figure S7). As PFB generates singlet oxygen with a ϕ_^_of 4%, it was important to show that photoactivation could be performed without provoking phototoxicity. In this endeavor cell viability MTT assays were conducted on cells incubated with PFB-Mito before and after activation. Both conditions showed no significant toxicity indicating no cytotoxicity of PFB-Mito as well as no phototoxicity upon photoactivation (Figure S8). Interestingly, PFB did not show more phototoxicity than the widely used Mito-Tracker™ Green after the same irradiation time at 488 nm (Figure S8).

To extend the versatility of PFB, a sulfonamide group and a Halo-tag group (Figure 2) were coupled to acid 4 to respectively target the endoplasmic reticulum (ER),^[39]^ and genetically encoded Halo-tagged proteins.^[40]^ The ER version of PFB, namely PFB-ER, was found to behave in a similar manner than for the mitochondrial version and sequential photoactivation could have been performed successfully (Figure 4C, figure S9), with a high selectivity toward the ER, (Pearson’s coefficient > 0.85, figure S10). Both PFB-Mito and PFB-ER were photoactivated in a fast manner and their photoactivated forms could be continuously imaged for up to 3 min after photoactivation with a slight decay due to progressive photobleaching (Figure S11).

PFB-Halo was then incubated with transfected cells expressing Halo-tagged proteins and was successfully photoactivated in a selective manner in various organelles, including nucleus (through the staining of chromatin), nucleoli and actin fibers (Figure 4C).

Overall, PFB was readily photoactivated in various organelles using both targeting moieties and genetically encoded proteins. PFB-Mito and PFB-ER provided images with high to excellent signal-to-noise ratios (SNR) after photoactivation up to 77 and with high SNR enhancement upon photoactivation (Figure 4C and S12, table S1). Although the signal of Halo-tagged fluorescent probes in transfected cells depends on the level of transfection frames and expression, the photoactivated PFB-Halo was also imaged with high SNR (Table S1).

### Live Super-Resolution Microscopy

To assess the efficiency of PFB in live single molecule localization microscopy, we used PFB-Halo in combination with Halo-tagged MAP4 protein to stain the thin microtubules’ cell cytoskeleton. The images were treated as single molecule localization microscopy (SMLM, figure 5A) using ThunderSTORM plugin,^[41]^ and as Super Resolution Radial Fluctuation (SRRF, figure 5B) method using an open-source plugin for ImageJ.^[42]^ Upon acquisition of SMLM movie and analysis of histogram of blinks over time, we noticed that PFB provided a high number of blinks over a short time frame (65,500 localizations within 1.4 s, Figure 5C) along with good localization precision of 27 nm (Figure 5C). These properties are consistent with our photoconvertible BODIPY based on directed photooxidation-induced conversion.^[29]^ Taking advantage of this feature, we successfully obtained super-resolution images of live cell’s microtubules (Figure 5A & B) by acquiring only 100 with an integration time of 14 ms per frame, which represents a total acquisition time of only 1.4 s. The two algorithms provided different, though complementary, super resolution images. Whereas thunderstorm plugin led to a higher resolution (67 nm, Figure 5D and S13), the SRRF plugin revealed more microtubules from less intense fluorescence signal fluctuations. With such a short time of acquisition, we challenged the system by performing sequential acquisitions of 1.4 s every 30 s. Surprisingly, PFB displayed a remarkably constant number of blinks over time with a stable localization precision (Figure 5C).

**Figure 5.**
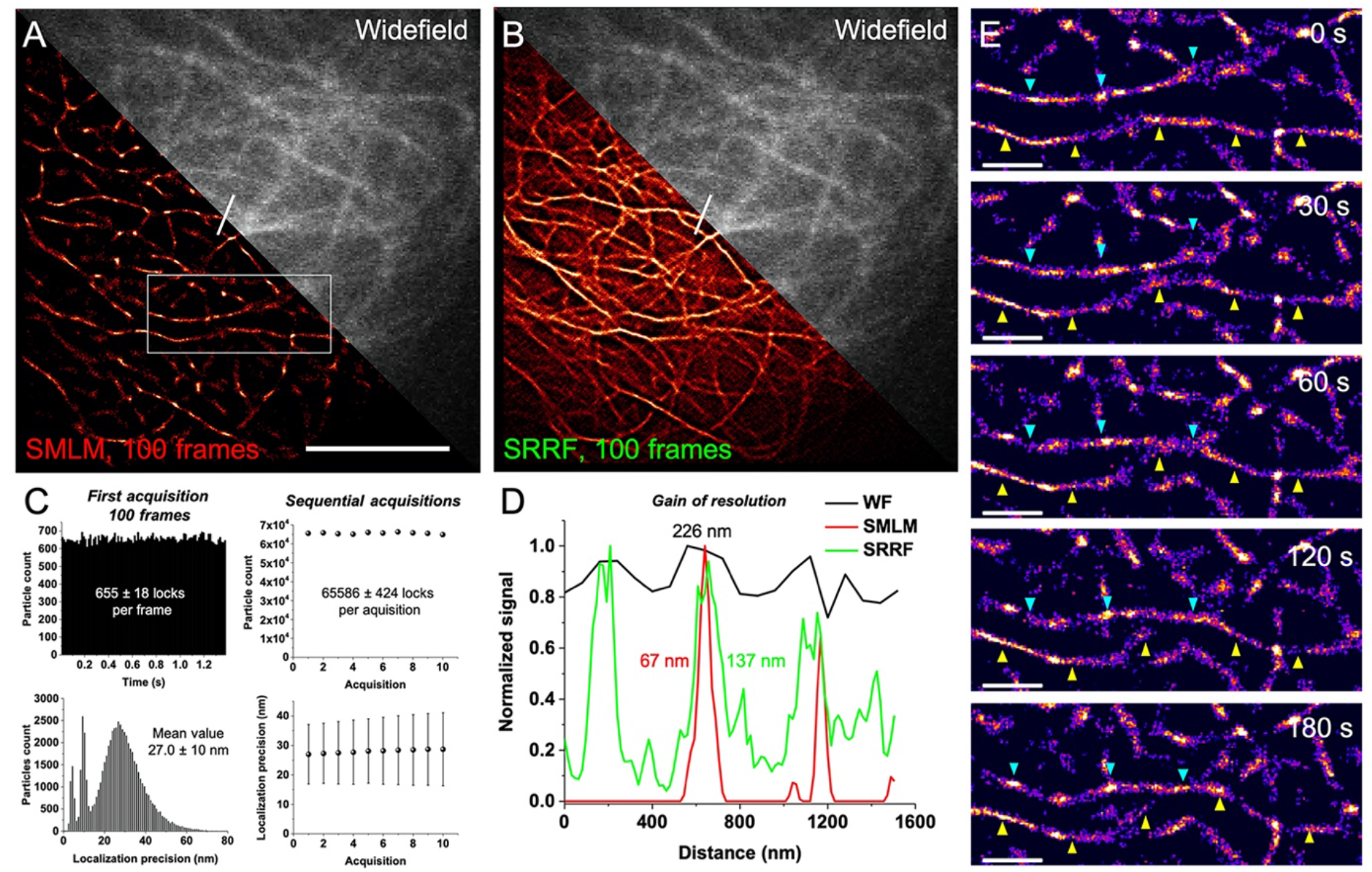
Live super-resolution imaging of HeLa cells’ microtubules using PFB-Halo (50 nM) and MAP4-Halo tagged protein. Comparison of the same field of view between SMLM and widefield images (A) and SRRF and widefield images (B). SMLM and SRRF images were reconstructed using only 100 frames acquisition for a total acquisition time of 1.4 s. Scale bar is 5 µm. (C) Histogram of number of blinks per frame and the localization precision for the first 100 frames acquisition showing the remarkable properties of PFB (left). Mean number of blinks and localization precision per acquisition along 10 successive acquisitions (100 frames each), enabling dynamic super-resolution imaging. (D) Plot profile corresponding to the white line in A and B and showing the gain of resolution provided by the two techniques compared to widefield microscopy. For the determination of resolution see figure S13. (E) Dynamic live SMLM corresponding to the white frame in A and showing movements of microtubules in high resolution. The cyan and yellow arrows indicate individual microtubules that seem to cross over time. Scale bar is 1 µm.

This protocol consisting in performing sequential acquisitions over the time thus enabled reconstructing super resolution movies showing dynamic of microtubules in live cells (Movie 1 for SMLM, movie 2 for SRRF). Using the dynamic SMLM sequence it was thus possible to track individual microtubule filaments with a high spatiotemporal resolution (Figure 5E).

## Conclusion

In conclusion, we showed that PFB bearing a furan (an Aromatic Singlet Oxygen Reactive Moiety) in β position of a BODIPY led to an efficient quenching of brightness. Upon irradiation at 488 nm, PFB generates ^1^O_2_ able to selectively protoxidize the furan giving rise to an impressive fluorescent enhancement of 93-fold at 510 nm. PFB was successfully targeted to various organelles using targeting moieties or Halo-tag and showed efficient sequential and selective photoactivation in live cells with high signal-to-noise ratios after photoactivation. In single molecule localization microscopy PFB displayed outstanding properties with an impressive number of blinks in short acquisition times allowing dynamic super resolution imaging in live cells. This work reaffirms the efficiency of directed photooxidation to develop advanced photomodulable fluorescent probes and extends the range of their applications.

## Supporting information

Moviie 2

## Supporting Information

Materials and methods, protocol and characterization of newly synthesized compounds, and supplementary figures can be found in the supplementary information.

**Movie 1.** Photoactivation of PFB-Mito using laser scanning confocal microscope.

**Movie 2.** Dynamic super resolution of microtubules using SMLM

**Movie 3.** Dynamic super resolution of microtubules using radial fluctuations.

## Acknowledgements

This work was founded by the Agence Nationale de la Recherche (ANR) BrightSwitch 19-CE29-0005-01, ANR 5D-SURE ANR-21-CE42-0015 and by the Ministère Français de l' Enseignement Supérieur, de la Recherche et de l' Innovation. The authors would like to thank Pr. Arnaud Gautier Dr. Halina Anton and Sophie Foppolo for providing the plasmids coding for Halo-tagged proteins and for mCherry. We also thank Dr. Andrey Klymchenko and Dr. Sophie Martin for granting us access to their equipment.

## Entry for the Table of Contents

PFB is an efficient, green-emitting, photoactivatable probe based on BODIPY. Upon irradiation with visible light (488 nm), directed photooxidation oxidizes the furanyl group and eliminates the quenching TICT mechanism, resulting in an impressive fluorescence enhancement at 510 nm (93-fold increase). PFB has been targeted to various cellular organelles and successfully used in photoactivation and live dynamic super-resolution microscopy. 

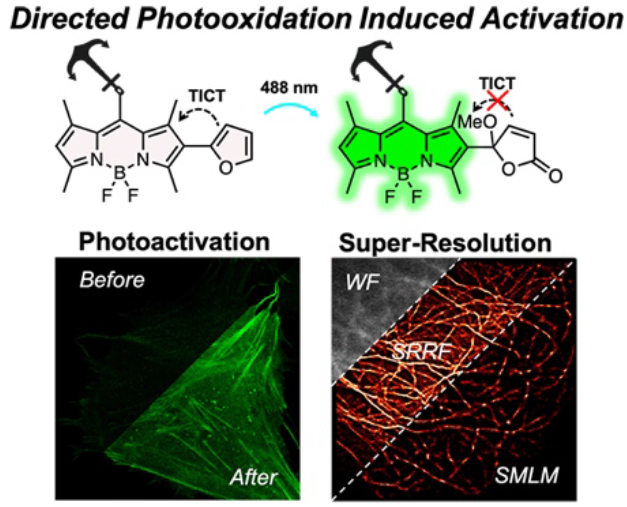

Institute and/or researcher Twitter usernames: @SaladinLazare, @vleberruyer, @Mayeul_Collot

## References

[1] K. A. Lukyanov, D. M. Chudakov, S. Lukyanov, V. V. Verkhusha, Nat Rev Mol Cell Biol 2005, 6, 885–890.

[2] Y. Zhang, F. M. Raymo, Methods Appl. Fluoresc. 2020, 8, 032002.

[3] S. J. Lord, N. R. Conley, H. D. Lee, R. Samuel, N. Liu, R. J. Twieg, W. E. Moerner, J. Am. Chem. Soc. 2008, 130, 9204–9205.

[4] M. S. Frei, P. Hoess, M. Lampe, B. Nijmeijer, M. Kueblbeck, J. Ellenberg, H. Wadepohl, J. Ries, S. Pitsch, L. Reymond, K. Johnsson, Nat Commun 2019, 10, 4580.

[5] X. Zhang, D. Guan, Y. Liu, J. Liu, K. Sun, S. Chen, Y. Zhang, B. Zhao, T. Zhai, Y. Zhang, F. Li, Q. Liu, Angewandte Chemie International Edition 2022, 61, e202211767.

[6] H. Li, J. C. Vaughan, Chem. Rev. 2018, 118, 9412–9454.

[7] K. Kikuchi, L. D. Adair, J. Lin, E. J. New, A. Kaur, Angewandte Chemie International Edition 2023, 62, e202204745.

[8] Y. Zhang, S. Swaminathan, S. Tang, J. Garcia-Amorós, M. Boulina, B. Captain, J. D. Baker, F. M. Raymo, J. Am. Chem. Soc. 2015, 137, 4709–4719.

[9] Y. Zhang, K.-H. Song, S. Tang, L. Ravelo, J. Cusido, C. Sun, H. F. Zhang, F. M. Raymo, J. Am. Chem. Soc. 2018, 140, 12741–12745.

[10] K. Nienhaus, G. U. Nienhaus, RSC Chem. Biol. 2021, 2, 796–814.

[11] V. Adam, R. Berardozzi, M. Byrdin, D. Bourgeois, Current Opinion in Chemical Biology 2014, 20, 92–102.

[12] V. N. Belov, C. A. Wurm, V. P. Boyarskiy, S. Jakobs, S. W. Hell, Angewandte Chemie International Edition 2010, 49, 3520–3523.

[13] L. M. Wysocki, J. B. Grimm, A. N. Tkachuk, T. A. Brown, E. Betzig, L. D. Lavis, Angewandte Chemie International Edition 2011, 50, 11206–11209.

[14] J. B. Grimm, T. Klein, B. G. Kopek, G. Shtengel, H. F. Hess, M. Sauer, L. D. Lavis, Angewandte Chemie International Edition 2016, 55, 1723–1727.

[15] R. Weber, S. Junek, A. Heckel, Chemistry – A European Journal n.d., n/a, e202300997.

[16] T. Kobayashi, T. Komatsu, M. Kamiya, C. Campos, M. González-Gaitán, T. Terai, K. Hanaoka, T. Nagano, Y. Urano, J. Am. Chem. Soc. 2012, 134, 11153–11160.

[17] H. He, Z. Ye, Y. Xiao, W. Yang, X. Qian, Y. Yang, Anal. Chem. 2018, 90, 2164–2169.

[18] J. B. Grimm, B. P. English, H. Choi, A. K. Muthusamy, B. P. Mehl, P. Dong, T. A. Brown, J. Lippincott-Schwartz, Z. Liu, T. Lionnet, L. D. Lavis, Nature Methods 2016, 13, 985–988.

[19] R. Lincoln, M. L. Bossi, M. Remmel, E. D’Este, A. N. Butkevich, S. W. Hell, Nat. Chem. 2022, 14, 1013–1020.

[20] I. Likhotkin, R. Lincoln, M. L. Bossi, A. N. Butkevich, S. W. Hell, J. Am. Chem. Soc. 2023, DOI 10.1021/jacs.2c12567.

[21] J. Tang, M. A. Robichaux, K.-L. Wu, J. Pei, N. T. Nguyen, Y. Zhou, T. G. Wensel, H. Xiao, J. Am. Chem. Soc. 2019, 141, 14699–14706.

[22] L. Wang, S. Wang, J. Tang, V. B. Espinoza, A. Loredo, Z. Tian, R. B. Weisman, H. Xiao, Chem. Sci. 2021, 12, 15572–15580.

[23] G. Ulrich, R. Ziessel, A. Harriman, Angewandte Chemie International Edition 2008, 47, 1184–1201.

[24] T. Kowada, H. Maeda, K. Kikuchi, Chem. Soc. Rev. 2015, 44, 4953–4972.

[25] Y. Jang, T.-I. Kim, H. Kim, Y. Choi, Y. Kim, ACS Appl. Bio Mater. 2019, 2, 2567–2572.

[26] C. S. Wijesooriya, J. A. Peterson, P. Shrestha, E. J. Gehrmann, A. H. Winter, E. A. Smith, Angewandte Chemie International Edition 2018, 57, 12685–12689.

[27] L. Saladin, V. Breton, O. Dal Pra, A. S. Klymchenko, L. Danglot, P. Didier, M. Collot, Angewandte Chemie International Edition 2023, 62, e202215085.

[28] L. Saladin, O. Dal Pra, A. S. Klymchenko, P. Didier, M. Collot, Chemistry – A European Journal 2023, n/a, e202203933.

[29] L. Saladin, V. Breton, V. Le Berruyer, P. Nazac, T. Lequeu, P. Didier, L. Danglot, M. Collot, J. Am. Chem. Soc. 2024, DOI 10.1021/jacs.4c05231.

[30] S. Dixit, T. Mahaddalkar, M. Lopus, N. Agarwal, Journal of Photochemistry and Photobiology A: Chemistry 2018, 353, 368–375.

[31] C.-W. Wan, A. Burghart, J. Chen, F. Bergström, L. B.-Å. Johansson, M. F. Wolford, T. G. Kim, M. R. Topp, R. M. Hochstrasser, K. Burgess, Chemistry – A European Journal 2003, 9, 4430–4441.

[32] M. K. Goshisht, N. Tripathi, G. K. Patra, M. Chaskar, Chem. Sci. 2023, DOI 10.1039/D3SC01036H.

[33] S. Sasaki, G. P. C. Drummen, G. Konishi, J. Mater. Chem. C 2016, 4, 2731–2743.

[34] R. Miao, J. Li, C. Wang, X. Jiang, Y. Gao, X. Liu, D. Wang, X. Li, X. Liu, Y. Fang, Advanced Science 2022, 9, 2104609.

[35] S. Sengupta, U. K. Pandey, Org. Biomol. Chem. 2018, 16, 2033–2038.

[36] T. Montagnon, D. Kalaitzakis, M. Triantafyllakis, M. Stratakis, G. Vassilikogiannakis, Chem. Commun. 2014, 50, 15480–15498.

[37] R. Ando, H. Hama, M. Yamamoto-Hino, H. Mizuno, A. Miyawaki, Proceedings of the National Academy of Sciences 2002, 99, 12651–12656.

[38] A. Acharya, A. M. Bogdanov, B. L. Grigorenko, K. B. Bravaya, A. V. Nemukhin, K. A. Lukyanov, A. I. Krylov, Chem. Rev. 2017, 117, 758–795.

[39] D. Singh, D. Rajput, S. Kanvah, Chem. Commun. 2022, 58, 2413–2429.

[40] G. V. Los, L. P. Encell, M. G. McDougall, D. D. Hartzell, N. Karassina, C. Zimprich, M. G. Wood, R. Learish, R. F. Ohana, M. Urh, D. Simpson, J. Mendez, K. Zimmerman, P. Otto, G. Vidugiris, J. Zhu, A. Darzins, D. H. Klaubert, R. F. Bulleit, K. V. Wood, ACS Chem. Biol. 2008, 3, 373–382.

[41] M. Ovesný, P. Křížek, J. Borkovec, Z. Švindrych, G. M. Hagen, Bioinformatics 2014, 30, 2389–2390.

[42] S. Culley, K. L. Tosheva, P. Matos Pereira, R. Henriques, The International Journal of Biochemistry & Cell Biology 2018, 101, 74–79.

