## Supplementary Information for "Targeted Photoactivatable Green-Emitting BODIPY Based on Directed Photooxidation Induced Activation and its Application to Live Dynamic Super-Resolution Microscopy"

### Supporting information

**Materials and Methods.** Materials. All starting materials for synthesis were purchased from Alfa Aesar (Haverhill, United States), Merck (Darmstadt, Germany), BLDpharm (Reinbek, Germany), or TCI Europe (Zwijndrecht, Belgium) and used as received unless stated otherwise. pcDNA3.1-mCherry was purchased from addgene (Watertown, USA). NMR spectra were recorded on a Bruker Avance III 400 MHz or 500 MHz spectrometers. Mass spectra were obtained using an Agilent Q-TOF 6520 mass spectrometer. Absorption spectra were recorded on a Cary 4000 spectrophotometer (Varian). Fluorescence spectra were recorded on a Fluoromax-4 (Jobin Yvon, Horiba) spectrofluorometer. Emission measurements were systematically done at 20˚C, unless indicated otherwise. All the spectra were corrected from the wavelength-dependent response of the detector. HPLC analyses were performed on Agilent 1260 Infinity II system equipped with Interchim PF5C18AQ-250/046 reversed phase column with gradual eluting from water to acetonitrile for 20 min at 1.5 mL/min. Protocols of synthesis of all new compounds are described below.

#### Protocol synthesis and characterization

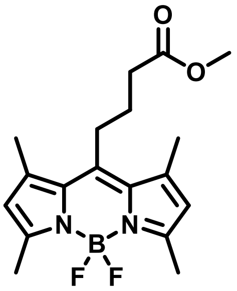


**1** was synthesized according to a published protocol.[1] To a solution of methyl 5-chloro-5-oxovalerate (3 mL, 22 mmol, 1 eq) in DCM (50 mL) was added dropwise 2,4-dimethylpyrrole (5 mL, 48 mmol, 2.2 eq) at 0°C. The solution was left to stir at RT for 30 minutes. TEA (8.9 mL, 65 mmol, 3eq) was added dropwise and the solution was left to stir at RT for 15 minutes. $BF_3.OEt_2$ (13.5 mL, 110 mmol, 5 eq) was added dropwise at 0°C and the solution was left to stir overnight at RT. The crude was filtered through silica using DCM, dry under reduced pressure and crystalized in EtOH to give **1** (3 g, 39%). $^1$H NMR (400 MHz, $CDCl_3$) δ 6.06 (s, 2H, ArH), 3.70 (s, 3H, OMe), 3.05 – 2.96 (m, 2H, $CH_2$ Alkyl), 2.55 – 2.47 (m, 8H, Dimethyl and $CH_2$ Alkyl), 2.43 (s, 6H, Dimethyl), 2.02 – 1.90 (m, 2H, $CH_2$ Alkyl).

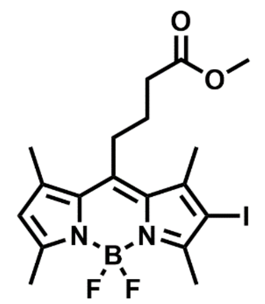

**2**. To a solution of **1** (200 mg, 574 µmol, 1 eq) in DMF (15 mL) was added ICl (30 µL, 574 µmol, 1eq) in MeOH (10 mL). The solution was left to stir at RT for 30 minutes. The solvents were removed under reduced pressure. The crude was purified by column chromatography on silica gel (DCM/Heptan : 40/60 to DCM 100%) to give **2** (200 mg, 73%). Rf = 0.6 (DCM). $^1$H NMR (400 MHz, $CDCl_3$) δ 6.13 (s, 1H, ArH), 3.70 (s, 3H, OMe), 3.07 – 2.99 (m, 2H, $CH_2$ Alkyl), 2.60 (s, 3H, Methyl), 2.54 – 2.49 (m, 5H, Methyl and $CH_2$ Alkyl), 2.47 (s, 3H, Methyl), 2.45 (s, 3H, Methyl), 2.00 – 1.88 (m, 2H, $CH_2$ Alkyl). $^{13}$C NMR (101 MHz, $CDCl_3$) δ 172.81, 156.51, 153.51, 144.77, 142.52, 140.17, 131.97, 130.97, 122.97, 122.95, 51.77, 34.03, 27.86, 26.70, 18.55, 16.63, 15.92, 14.65. HRMS (ESI+), calculated for $C_{18}H_{22}BFIN_2O_2$ $[M-F]^+$ 455.0803, found 455.0803.

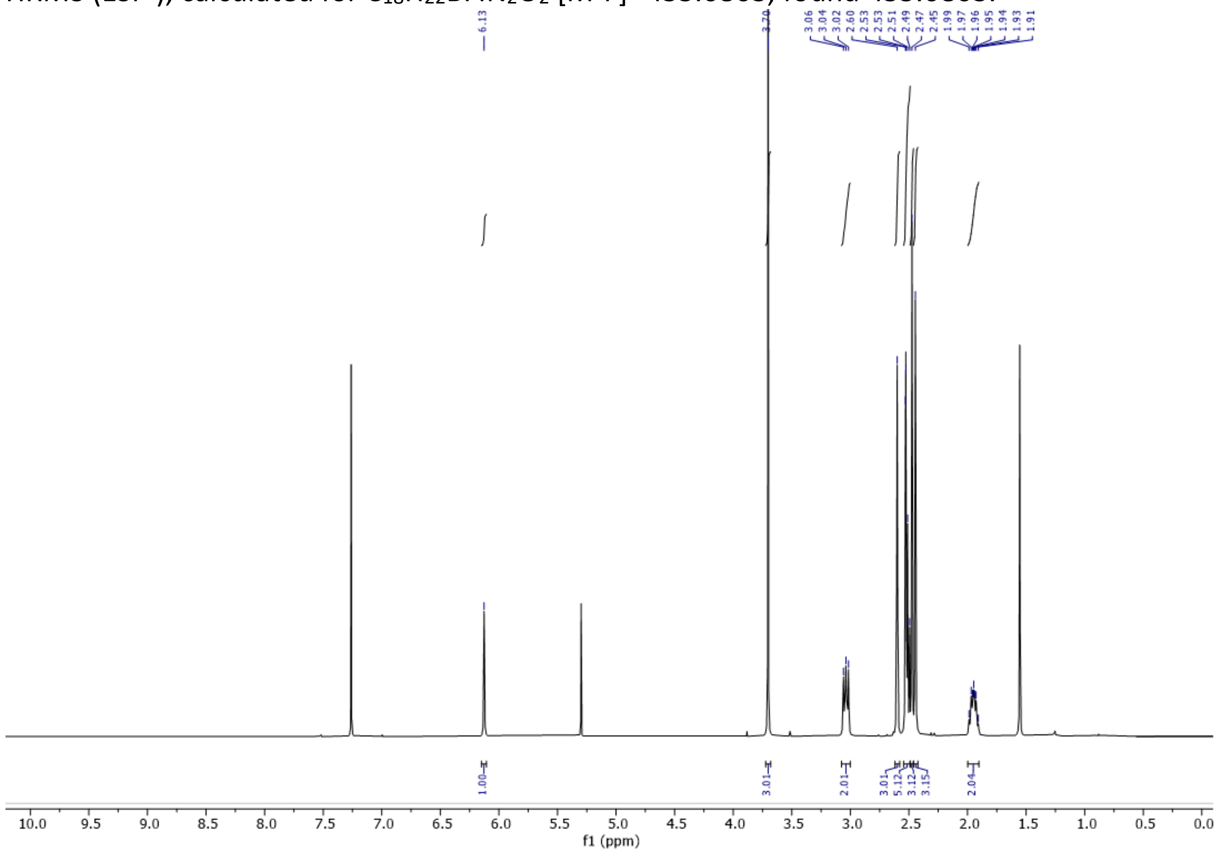


$^1$H NMR spectrum of **2**

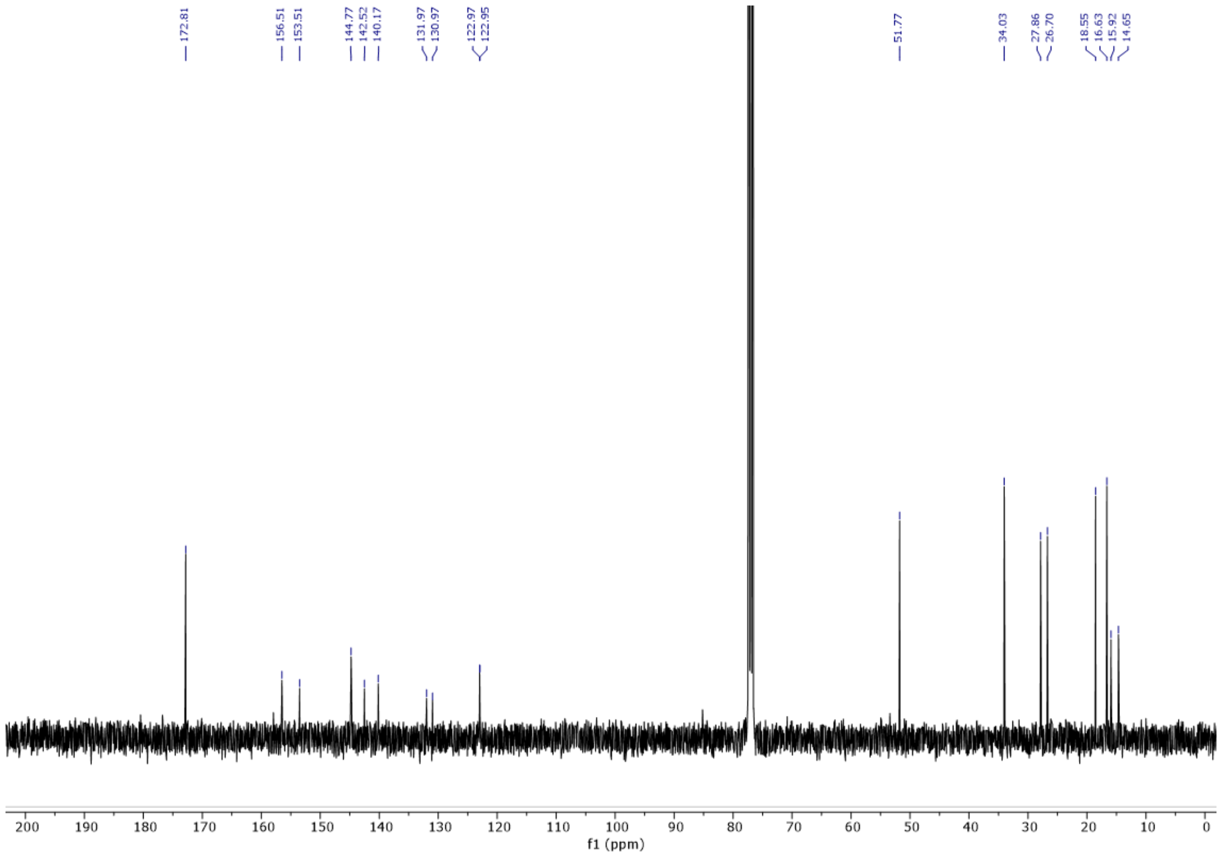


$^{13}C$ NMR spectrum of **2**

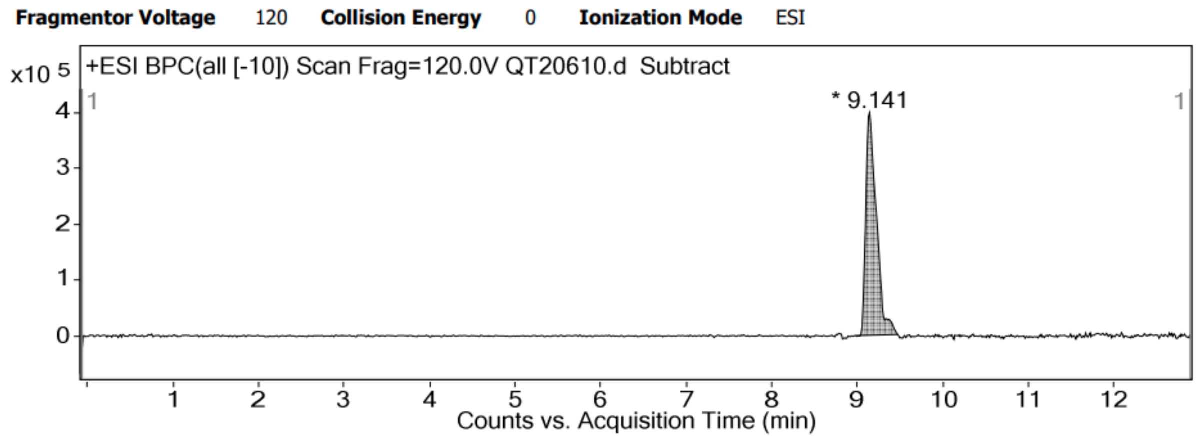


HPLC trace of **2**

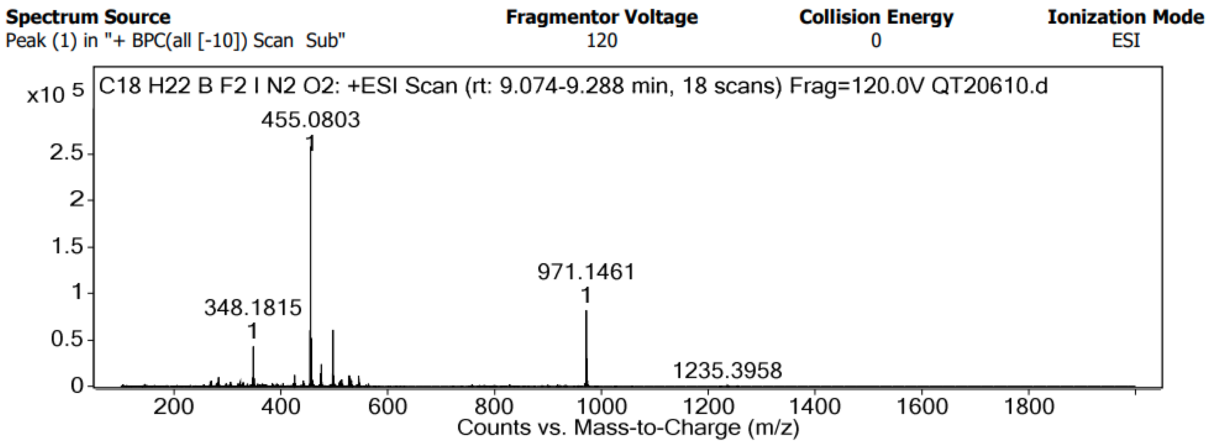

HRMS spectrum of **2**

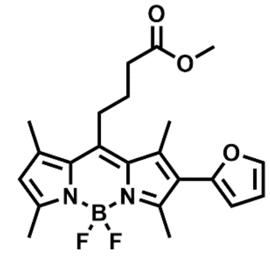

**3**. To a solution of **2** (100 mg, 232 µmol, 1 eq) in Dioxane (3 mL) and $Na_2CO_3$ aq (10%, 2 mL) was added 2- furanboronic acid (28 mg, 255 µmol, 1.1 eq) and $Pd(PPh_3)_4$ (27 mg, 0.1 eq). The reaction mixture was heated at 100°C fo 2 hours. The solvents were removed under reduced pressure. The crude was purified by column chromatography on silica gel (DCM/Heptan : 70/30) to give **3** (90 mg, 94%). $^1$H NMR (400 MHz, $CDCl_3$) δ 7.54 – 7.49 (m, 1H, Furan), 6.50 (dd, J = 3.3, 1.9 Hz, 1H, Furan), 6.31 (dd, J = 3.3, 0.8 Hz, 1H, Furan), 6.11 (s, 1H, ArH), 3.70 (s, 3H, OMe), 3.09 (m, 2H, Alkyl), 2.60 (s, 3H, Methyl), 2.55 – 2.50 (m, 5H Methyl and Alkyl), 2.48 (s, 3H, Methyl), 2.46 (s, 3H, Methyl), 2.00 (m, 2H, Alkyl). $^{13}$C NMR (126 MHz, $CDCl_3$) δ 172.88, 155.42, 153.02, 148.02, 145.53, 142.04, 141.29, 136.70, 132.15, 130.86, 122.47, 110.87, 109.31, 51.74, 34.10, 27.61, 26.82, 16.56, 14.58, 14.26, 13.57. HRMS (ESI+), calculated for $C_{22}H_{25}BFN_2O_3$ $[M-F]^+$ 395.1942, found 395.1976.

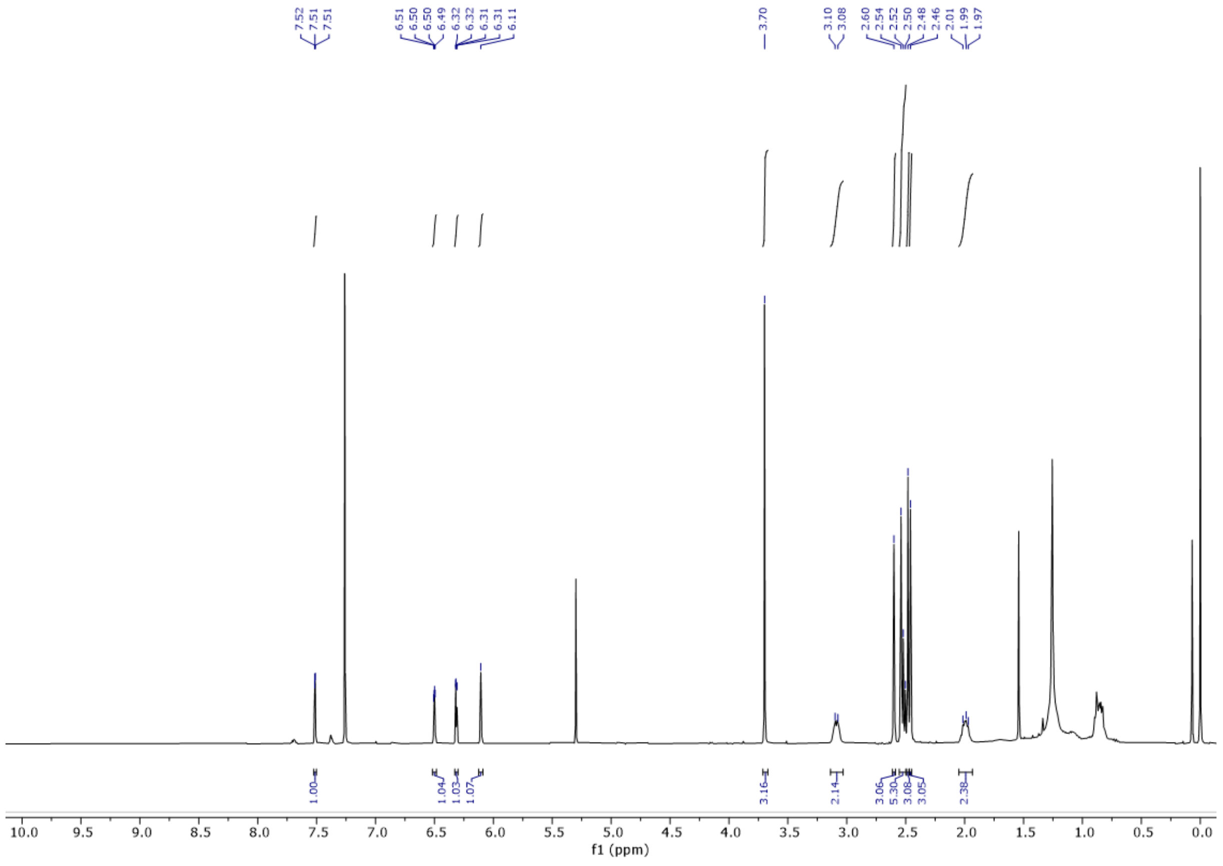


$^1$H NMR spectrum of **3**

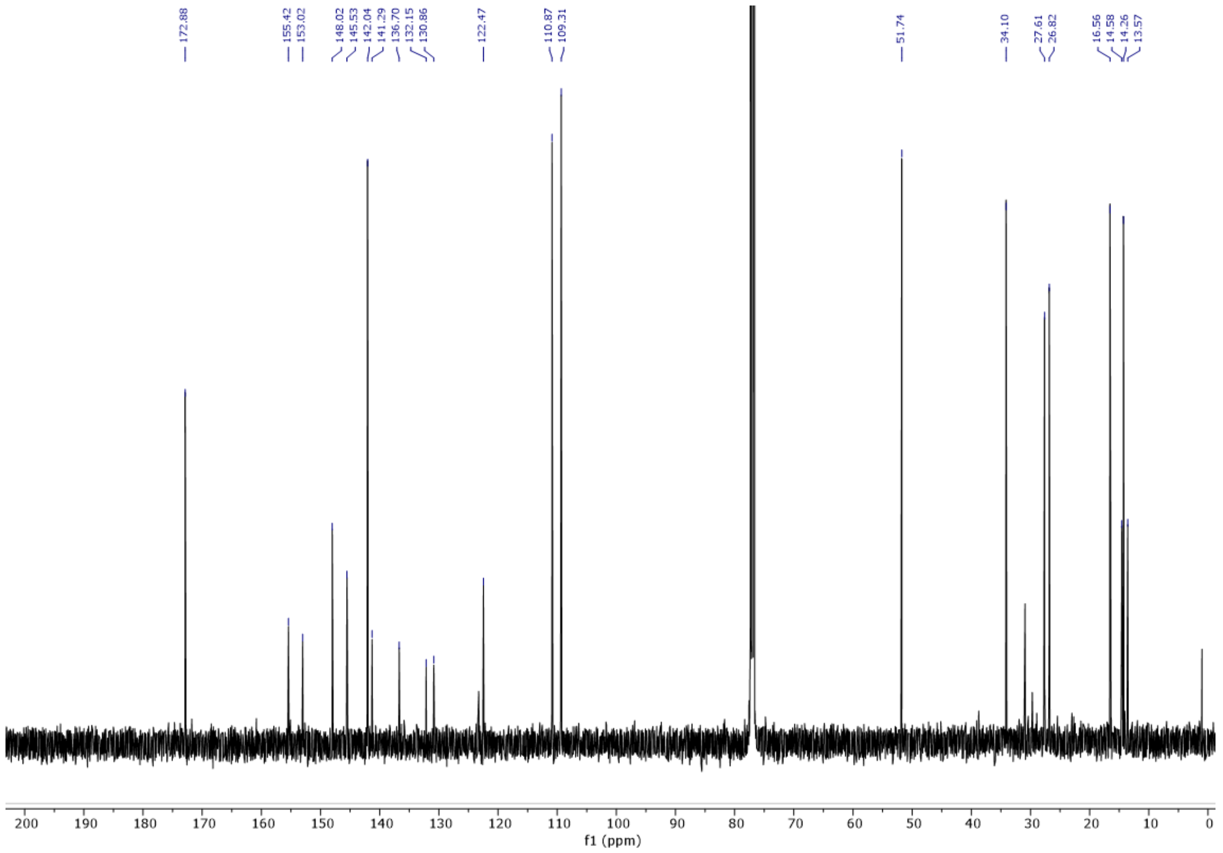


$^{13}$C NMR spectrum of **3**

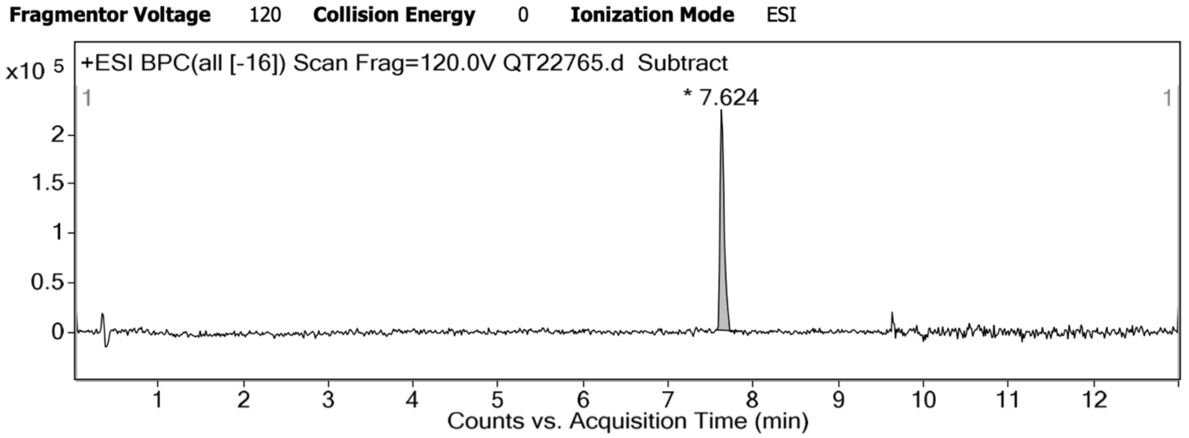


HPLC trace of **3**

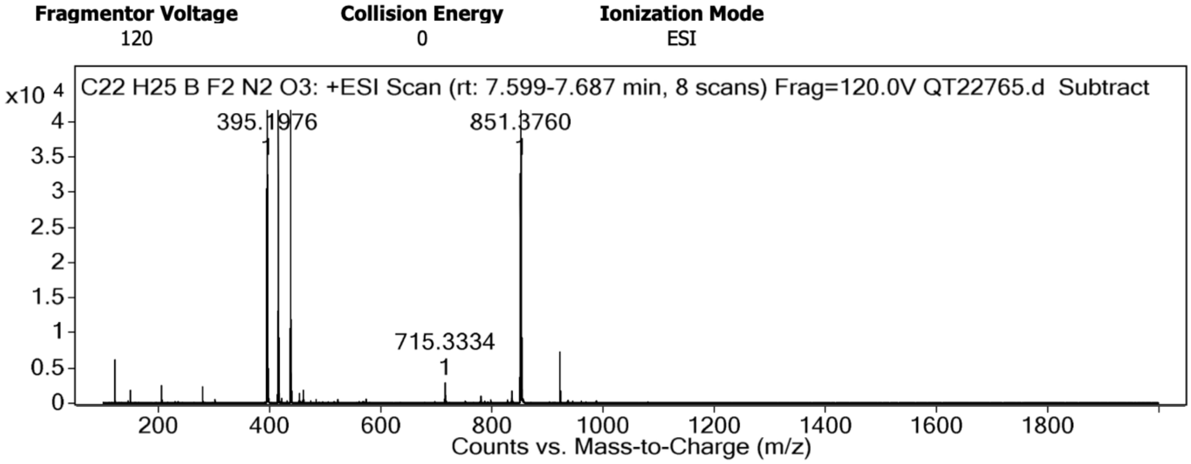


HRMS spectrum of **3**

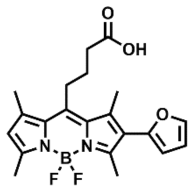


**4**. To a solution of **3** (200 mg, 483 µmol, 1 eq) in THF (20 ml) was added 4.8 ml of a solution of NaOH (0.1 M). The mixture was left to stir for 2 hours was was then quenched with HCl 1M. The solvent were removed under reduced pressure and the crude was purified by column chromatography on silica gel (DCM/MeOH : 9/1) to give **4** (120 mg, 62%). Rf = 0.57 (DCM/MeOH : 9/1). $^{1}$H NMR (400 MHz, $CDCl_3$) δ 7.51 (d, J = 1.8 Hz, 1H, ArH Furan), 6.49 (dd, J = 3.3, 1.9 Hz, 1H, ArH Furan), 6.31 (d, J = 3.3 Hz, 1H, ArH Furan), 6.10 (s, 1H, ArH BODIPY), 3.10 (d, J = 9.8 Hz, 2H, Alkyl), 2.60 (s, 3H, Methyl), 2.55 (m, 5H, Methyl and Alkyl), 2.47 (s, 3H, Methyl), 2.45 (s, 3H, Methyl), 2.01 (d, J = 9.5 Hz, 2H, Alkyl). $^{13}$C NMR (101 MHz, $CDCl_3$) δ 177.83, 155.52, 153.15, 147.97, 145.24, 142.06, 141.22, 136.67, 132.15, 130.86, 123.33, 122.52, 110.87, 109.34, 33.86, 27.50, 26.49, 16.53, 14.56, 14.24, 13.56. HRMS (ESI+), calculated for $C_{21}H_{23}BFN_2O_3$ $[M-F]^+$ 381.1786, found 381.1788.

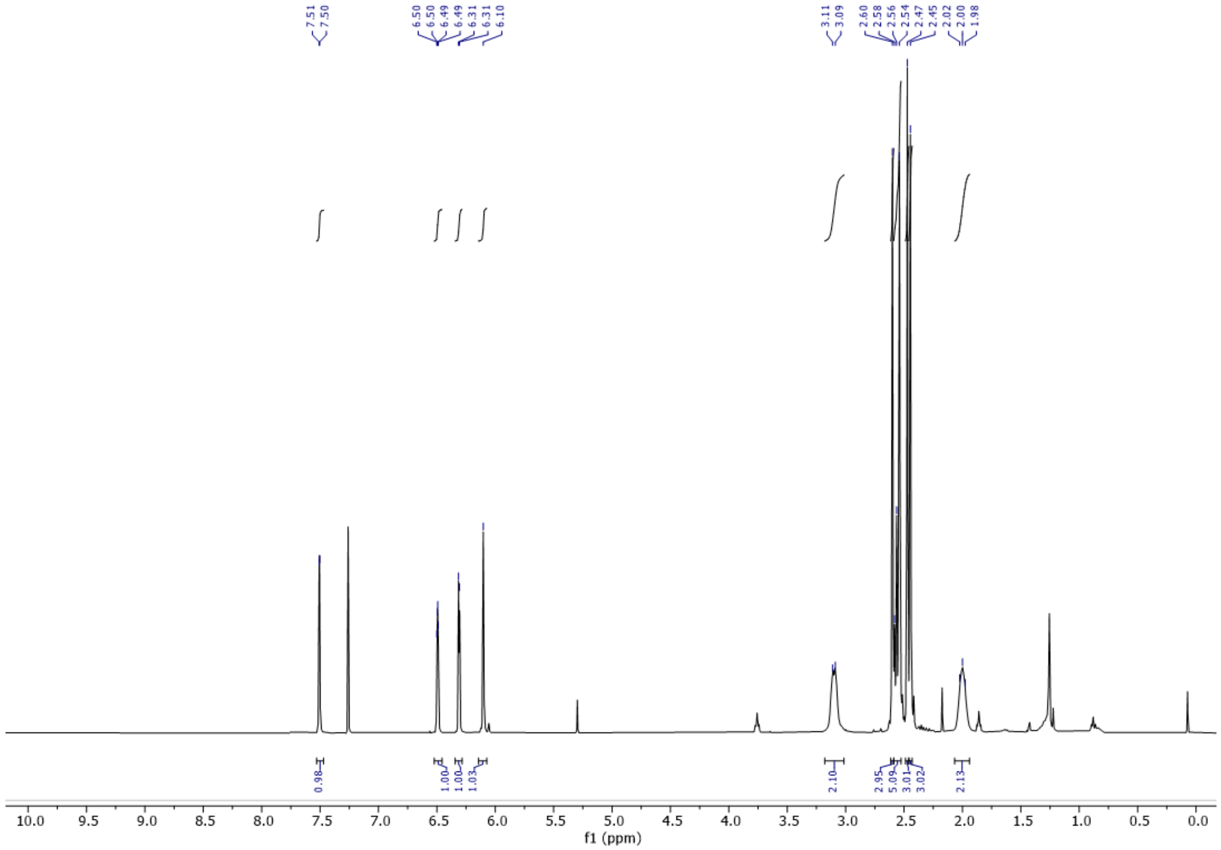


$^{1}$H NMR spectrum of 4

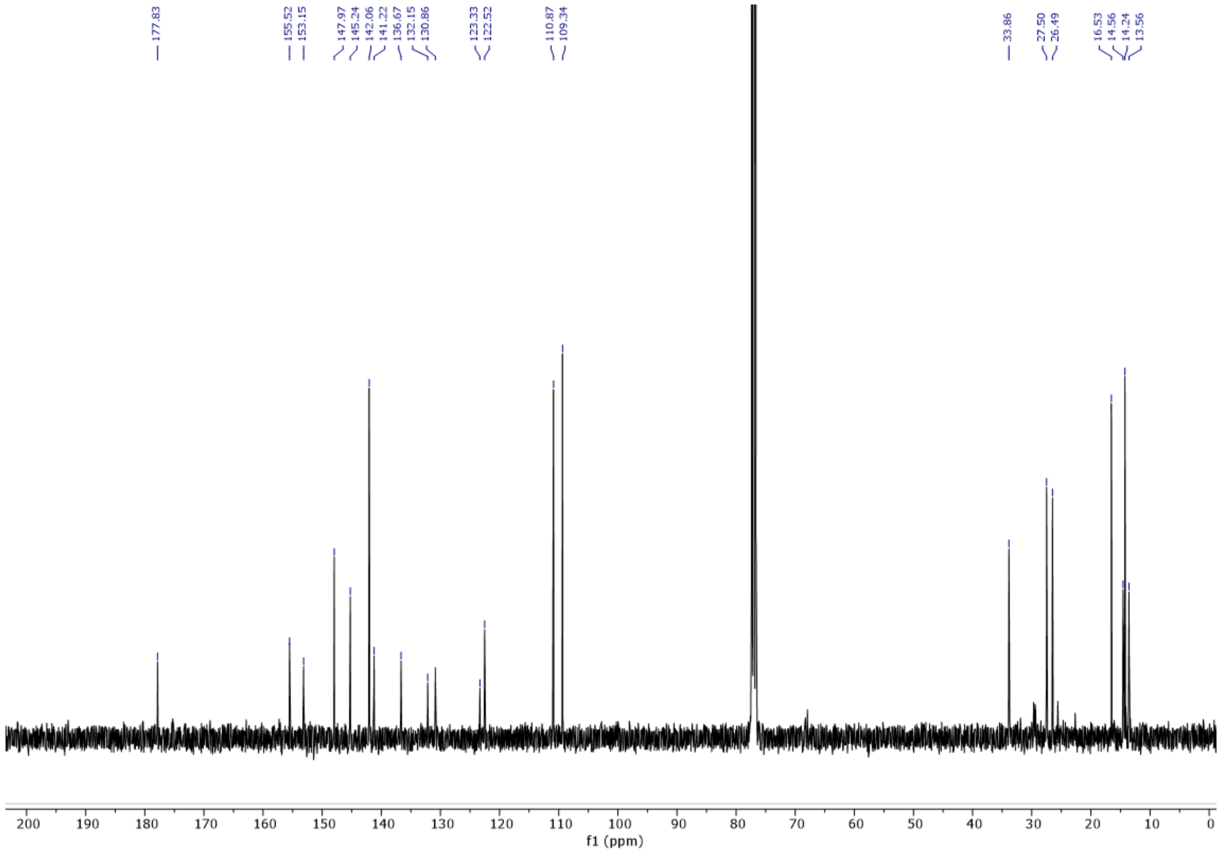


$^{13}$C NMR spectrum of **4**

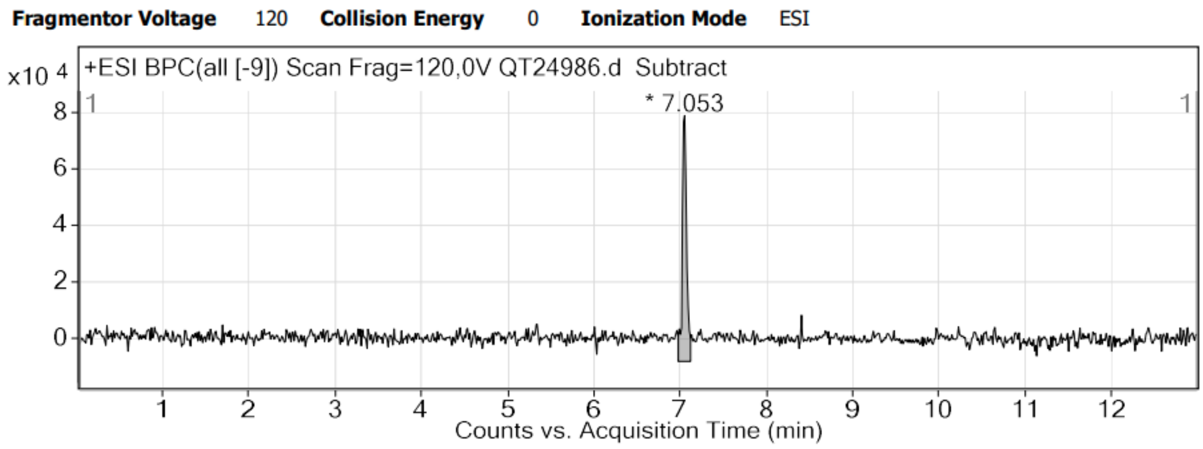


HPLC trace of **4**

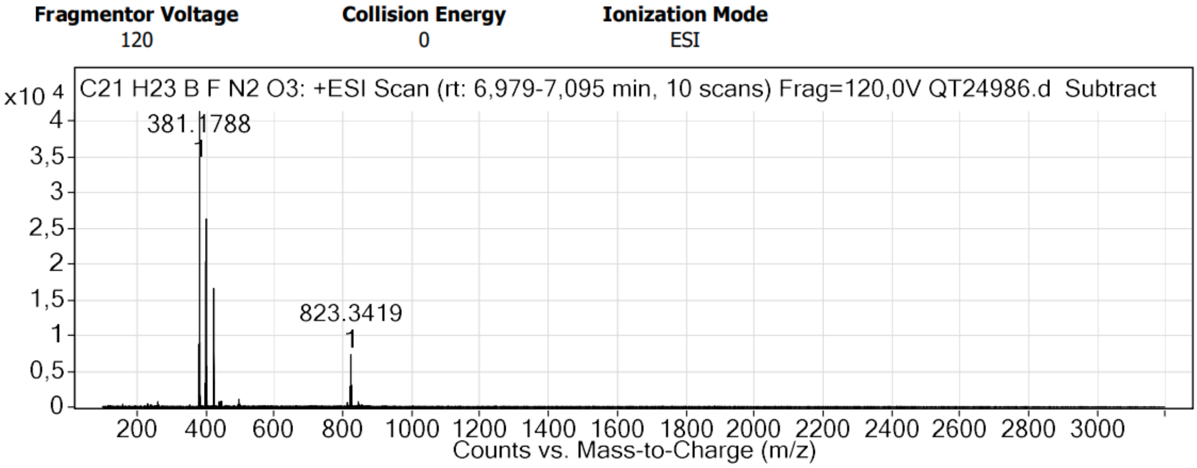


HMRS spectrum of **4**

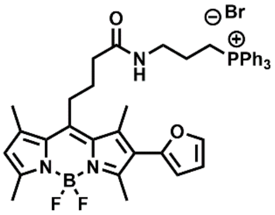

**PFB-Mito**. To a solution of 4 (80 mg, 193 µmol, 1 eq) in THF (10 mL) was added NaOH 0.1M (1.93 mL, 193 µmol, 1 eq). The reaction was left to stir one hour. The solvents were removed under reduced pressure. The crude was dissolved in DMF (3 mL), then HATU (85 mg, 223 µmol, 1.2 eq), **7** (134 mg, 279 µmol, 1.5 eq) and DIEA (97 µL, 558 µmol, 3 eq) were added. The reaction was left to stir for 10 minutes. The solvent were removed under reduced pressure and the crude was purified by column chromatography on silica gel (DCM/MeOH : 98/2) to give **PFB-Mito** (20 mg, 14%). Rf = 0.55 (DCM/MeOH : 9/1). $^{1}$H NMR (400 MHz, $CDCl_3$) δ 7.79 – 7.54 (m, 15H, $PPh_3$), 7.48 (dd, J = 2.0, 0.7 Hz, 1H, Furan), 6.87 (t, J = 6.1 Hz, 1H, NH Amide), 6.48 (dd, J = 3.3, 1.9 Hz, 1H, Furan), 6.29 (dd, J = 3.3, 0.8 Hz, 1H, Furan), 6.07 (s, 1H, BODIPY), 3.41 (p, J = 6.7, 6.0 Hz, 2H, Alkyl), 3.18 (ddt, J = 13.1, 8.2, 4.9 Hz, 2H, Alkyl), 2.97 (t, J = 8.7 Hz, 2H, Alkyl), 2.90 (s, 3H, Methyl), 2.58 (s, 3H, Methyl), 2.51 (s, 3H, Methyl), 2.40 (s, 8H, Methyl and Alkyl), 1.91 – 1.80 (m, 4H, Alkyl). $^{13}$C NMR (101 MHz, $CDCl_3$) δ 173.15, 155.26, 152.62, 148.18, 146.39, 142.22, 142.02, 137.29, 135.50, 133.57, 133.47, 133.37, 133.27, 130.92, 130.87, 130.80, 130.75, 130.67, 118.22, 117.86, 117.37, 117.00, 111.05, 109.36, 36.11, 27.93, 27.80, 22.61, 16.64, 14.66, 14.30, 13.68. HRMS (ESI+), calculated for $C_{42}H_{44}BF_2N_3O_2P$ $[M]^+$ 702.3227, found 702.3244.

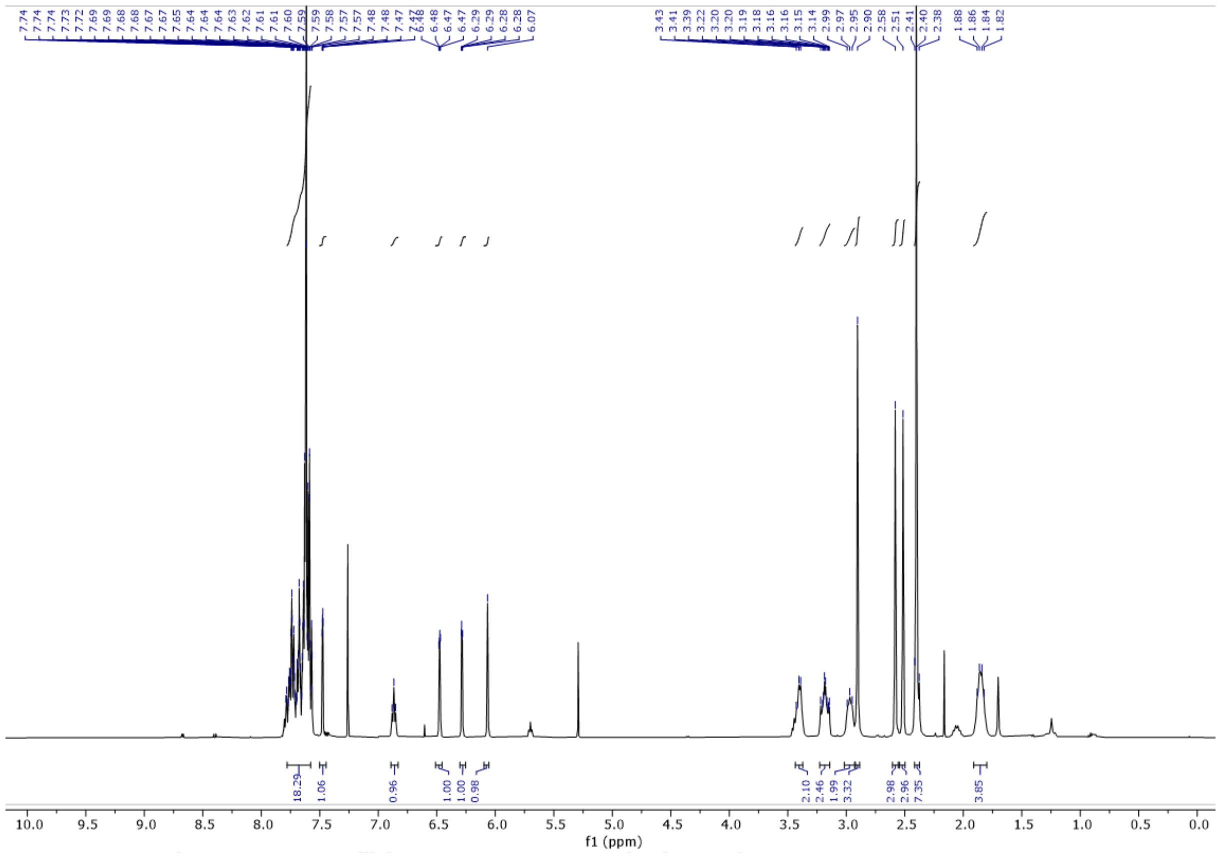

$^1$H NMR spectrum of **PFB-Mito**

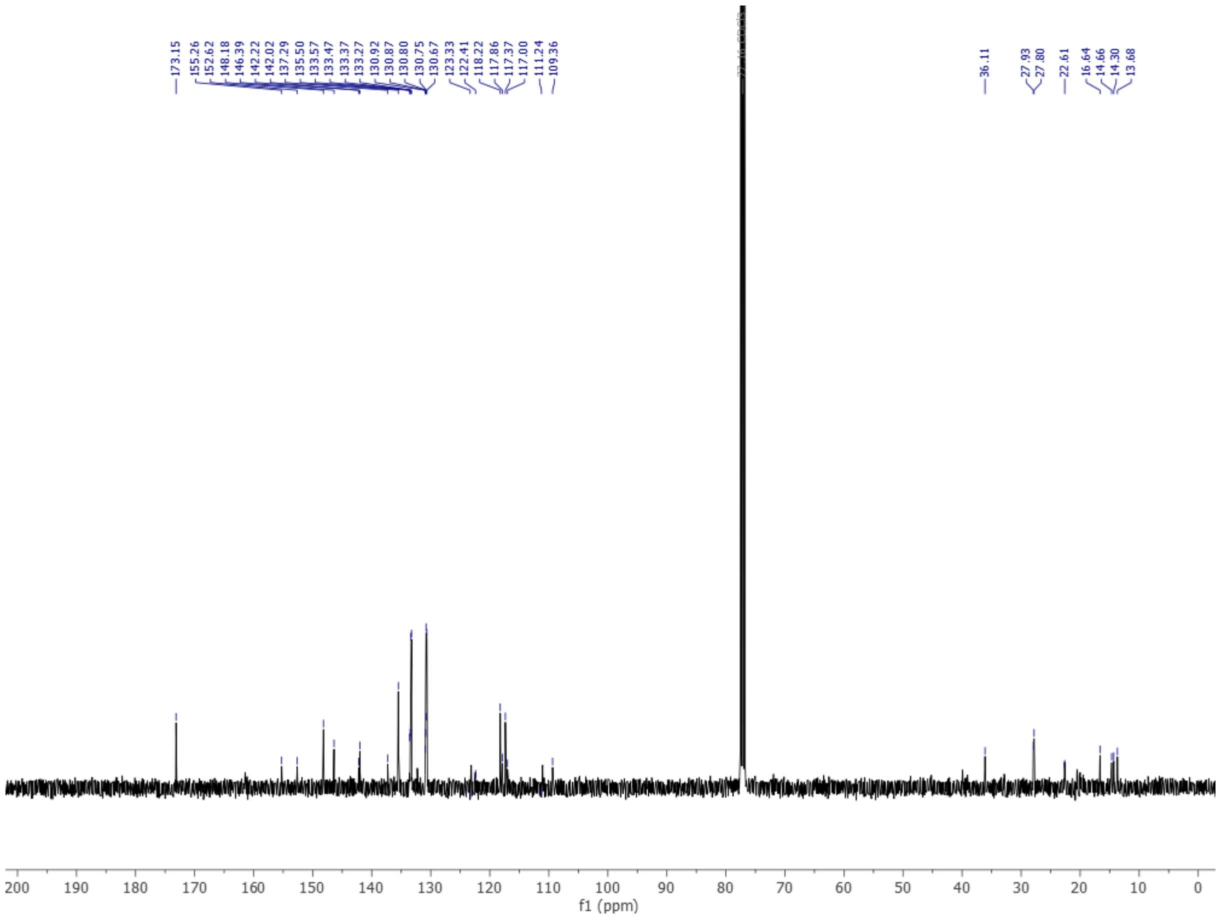


$^{13}$C NMR spectrum of **PFB-Mito**

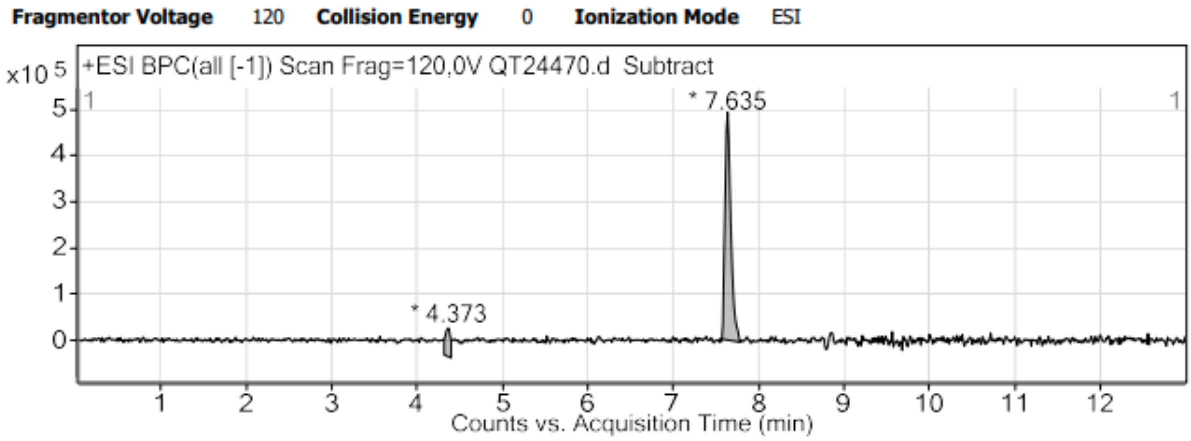


HPLC trace of **PFB-Mito**

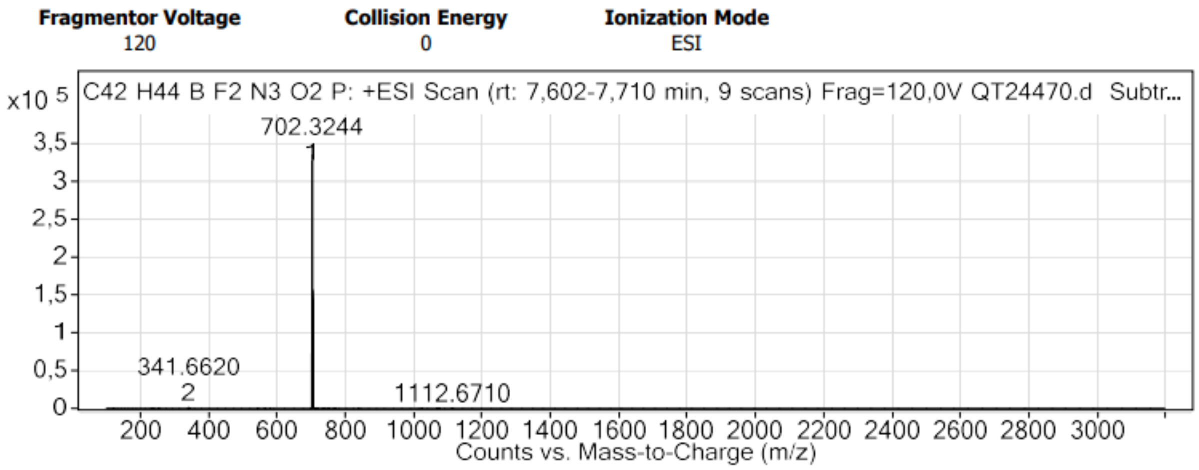


HRMS spectrum of **PFB-Mito**

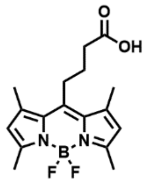

**6** was synthesized according to a published protocol.[1] To a solution of **1** (1.12 g, 3.23 mmol, 1 eq) in THF (60 mL) was added 32 mL of a solution of NaOH (0.1M). The mixture was left to stir for 15 minutes and was then quenched with HCl 6M. The precipitate was filtered then washed with water to give **8** (934 mg, 87%). $^1$H NMR (400 MHz, $CDCl_3$) δ 7.81 – 7.72 (m, 3H, $PPh_3$), 7.69 – 7.55 (m, 12H, $PPh_3$), 6.85 (t, J = 6.1 Hz, 1H, NH Amide), 6.02 (s, 2H, BODIPY), 3.72 – 3.64 (m, 4H, Alkyl), 3.40 (q, *J* = 5.4 Hz, 2H, Alkyl), 3.20 – 3.12 (m, 4H), 2.94 – 2.88 (m, 2H, Alkyl), 2.48 (s, 6H, Methyl), 2.36 (s, 6H, Methyl). $^{13}$C NMR (101 MHz, $CDCl_3$) δ residual solvent DMF, 173.29, 162.84, 153.95, 145.76, 141.19, 135.51, 133.39, 133.29, 131.56, 130.82, 130.70, 121.80, 118.19, 117.33, 55.75, 43.70, 36.20, 28.04, 27.64, 22.55, 18.59, 17.12, 16.37, 14.54, 12.77.

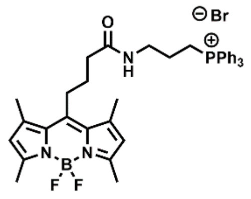

**MB-Mito**. To a solution of 6 (50 mg, 150 µmol, 1eq) in DMF (3 mL) was added HATU (68 mg, 180 µmol, 1.2 eq), **5** (108 mg, 224 µmol, 1.5 eq) and DIEA (78 µL, 449 µmol, 3 eq) were added. The reaction was left to stir for one hour. The solvent were removed under reduced pressure and the crude was purified by column chromatography on silica gel (DCM/MeOH : 9/1) to give **MB-Mito** (50 mg, 47%). Rf = 0.3 (DCM/MeOH : 9/1). $^1$H NMR (400 MHz, $CDCl_3$) δ 7.81 – 7.72 (m, 3H, $PPh_3$), 7.69 – 7.55 (m, 12H, $PPh_3$), 6.85 (t, J = 6.1 Hz, 1H, NH Amide), 6.02 (s, 2H, BODIPY), 3.72 – 3.64 (m, 4H, Alkyl), 3.40 (q, J = 5.4 Hz, 2H, Alkyl), 3.20 – 3.12 (m, 4H), 2.94 – 2.88 (m, 2H, Alkyl), 2.48 (s, 6H, Methyl), 2.36 (s, 6H, Methyl). $^{13}$C NMR (101 MHz, $CDCl_3$) δ 173.29, 162.84, 153.95, 145.76, 141.19, 135.51, 133.39, 133.29, 131.56, 130.82, 130.70, 121.80, 118.19, 117.33, 55.75, 43.70, 36.20, 28.04, 27.64, 22.55, 18.59, 17.12, 16.37, 14.54, 12.77. HRMS (ESI+), calculated for $C_{38}H_{42}BF_2N_3OP$ $[M]^+$ 636.3121, found 636.3137.

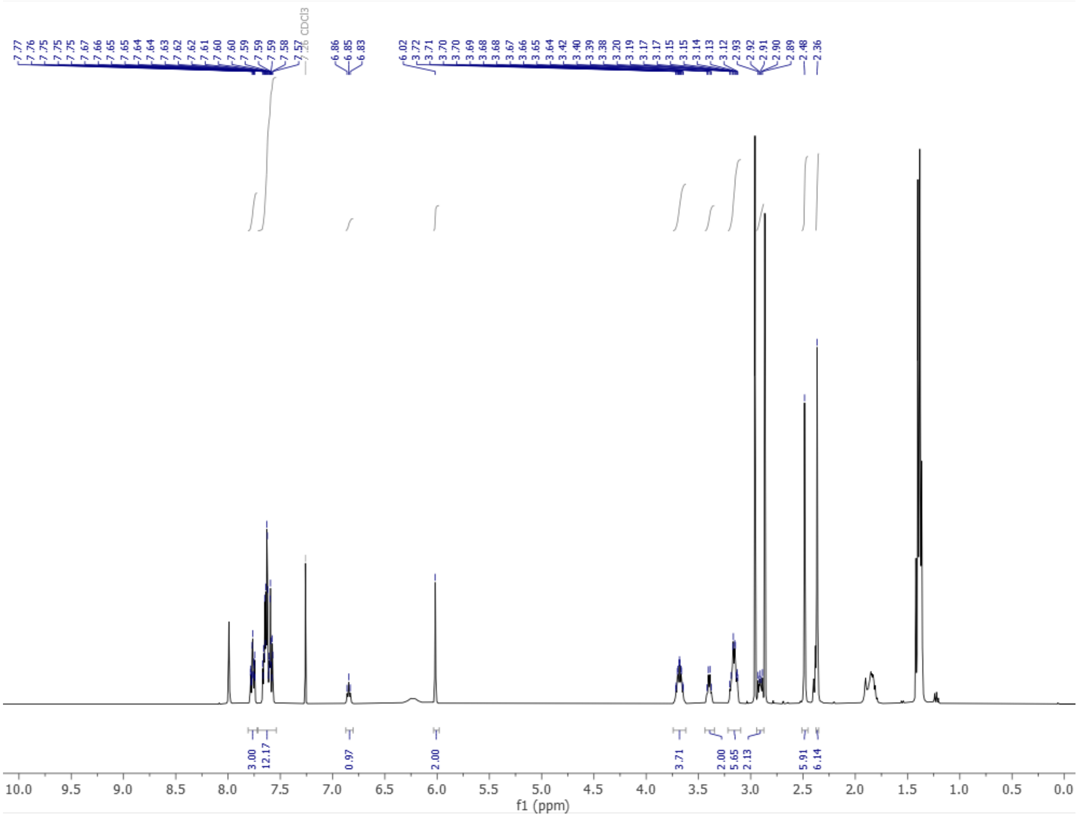


$^{1}H$ NMR spectrum of **MB-Mito**

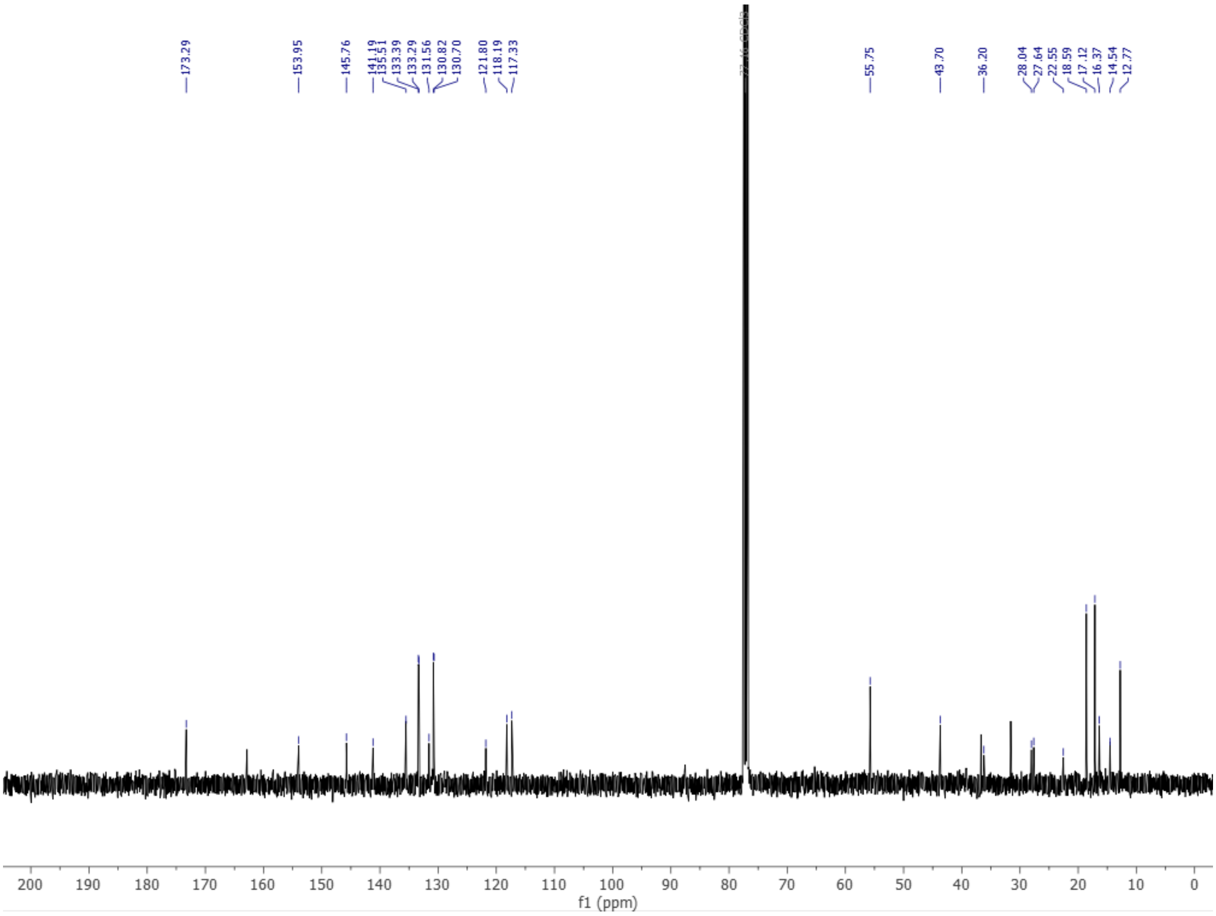


$^{13}C$ NMR spectrum of **MB-Mito**

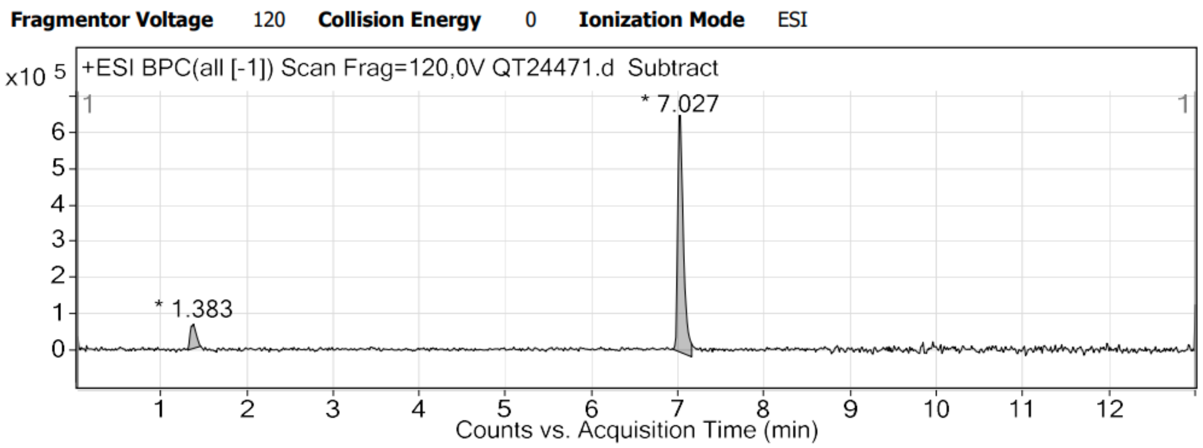


HPLC trace of **MB-Mito**

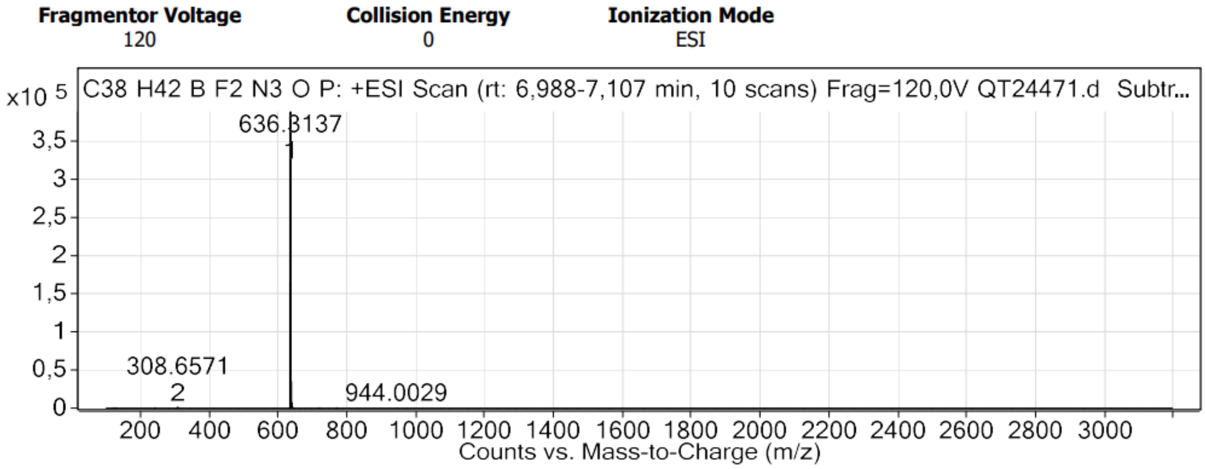


HRMS spectrum of **MB-Mito**

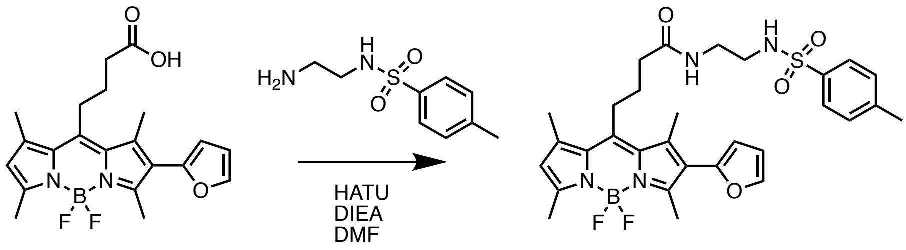


**4** **PFB-ER**

**PFB-ER**. To a solution of **4** (10 mg, 25 µmol, 1 eq) in DMF (2 mL) was added N-(2-((2-Aminoethyl)amino)ethyl)-4-methylbenzenesulfonamide (6.4 mg, 30 µmol, 1.2 eq), HATU (12.5 mg, 32 µmol, 1.3 eq) and DIEA (9 µL, 50 µmol, 2 eq). The solution was left to stir for 3 hours. The crude was concentrated under reduced pressure and purified by column chromatography on silica gel (DCM/MeOH: 9/1) to obtain **4** (7 mg, 47%). Rf = 0.47 (DCM/MeOH : 9/1).$^{1}$H NMR (500 MHz, $CDCl_3$) δ 7.70 (d, J = 8.0 Hz, 2H, ArH Tosyl), 7.50 (d, J = 1.9 Hz, 1H, ArH Furan), 7.27 (d, J = 8.3Hz, 2H, ArH Tosyl), 6.49 (dd, J = 3.2, 1.9 Hz, 1H, ArH Furan), 6.30 (d, *J* = 3.3 Hz, 1H, ArH Furan), 6.22 (br, 1H, Sulfonamide), 6.08 (s, 1H, ArH BODIPY), 5.33 (br, 1H, Amide), 3.34 (q,

J = 5.5 Hz, 2H, Alkyl), 3.04 (q, J = 5.7 Hz, 4H), 2.59 (s, 3H, Methyl), 2.53 (s, 3H, Methyl), 2.44 (s, 3H, Methyl), 2.42 (s, 3H, Methyl), 2.38 (s, 3H, Methyl), 2.34 (t, J = 7.1 Hz, 2H, Alkyl), 2.00 – 1.91 (m, 2H, Alkyl). $^{13}C$ NMR (126 MHz, $CDCl_3$) δ 172.78, 155.48, 152.97, 148.16, 146.01, 143.99, 142.14, 141.69, 136.93, 136.64, 132.35, 131.02, 130.00, 127.13, 122.59, 111.04, 109.45, 43.23, 39.59, 36.23, 29.84, 27.77, 27.39, 21.60, 16.75, 14.71, 14.47, 14.25, 13.72. HRMS (ESI+) calculated for $C_{30}H_{35}BFN_4O_4S$ $[M-F]^+$ 577.2456, found 577.2465

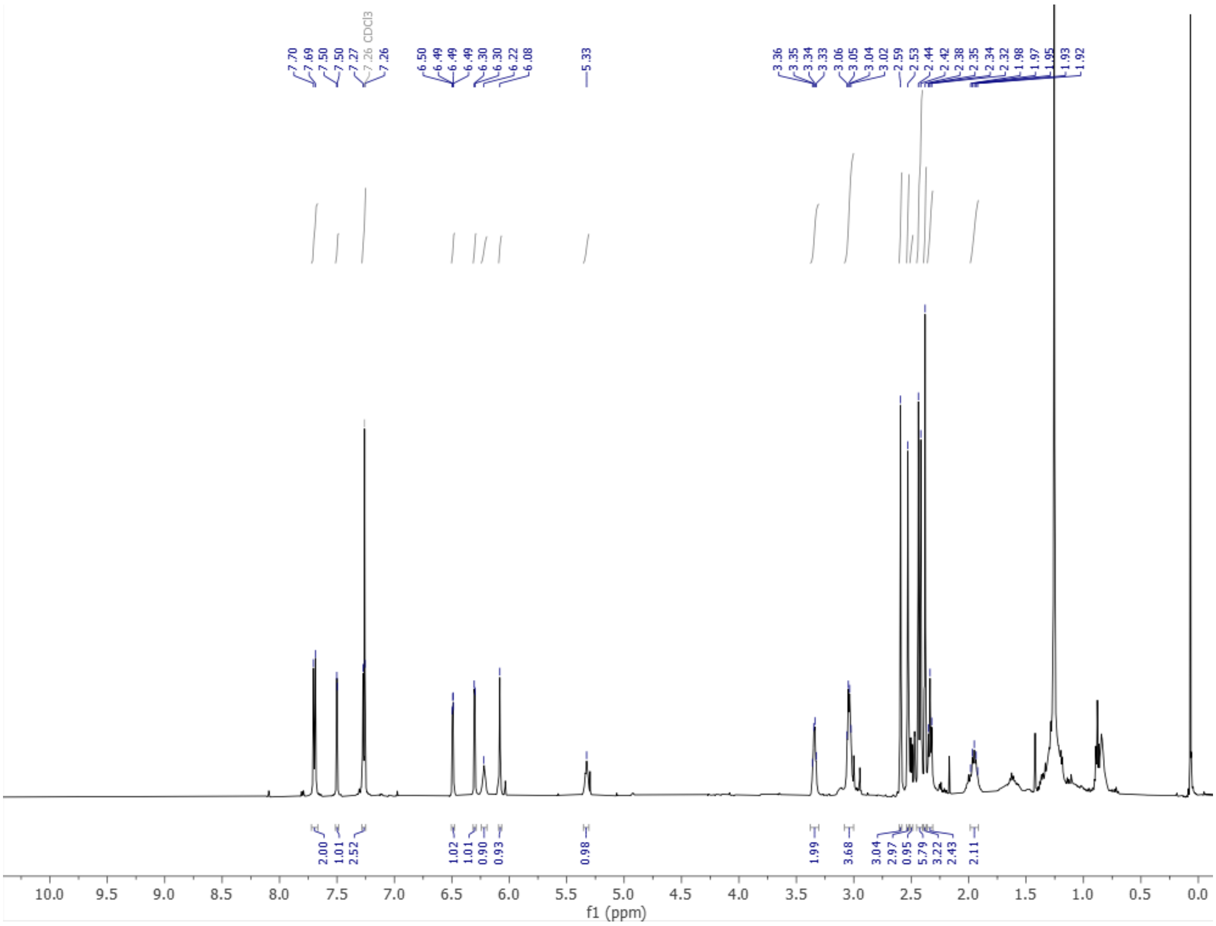

$^1H$ NMR spectrum of **PFB-ER**

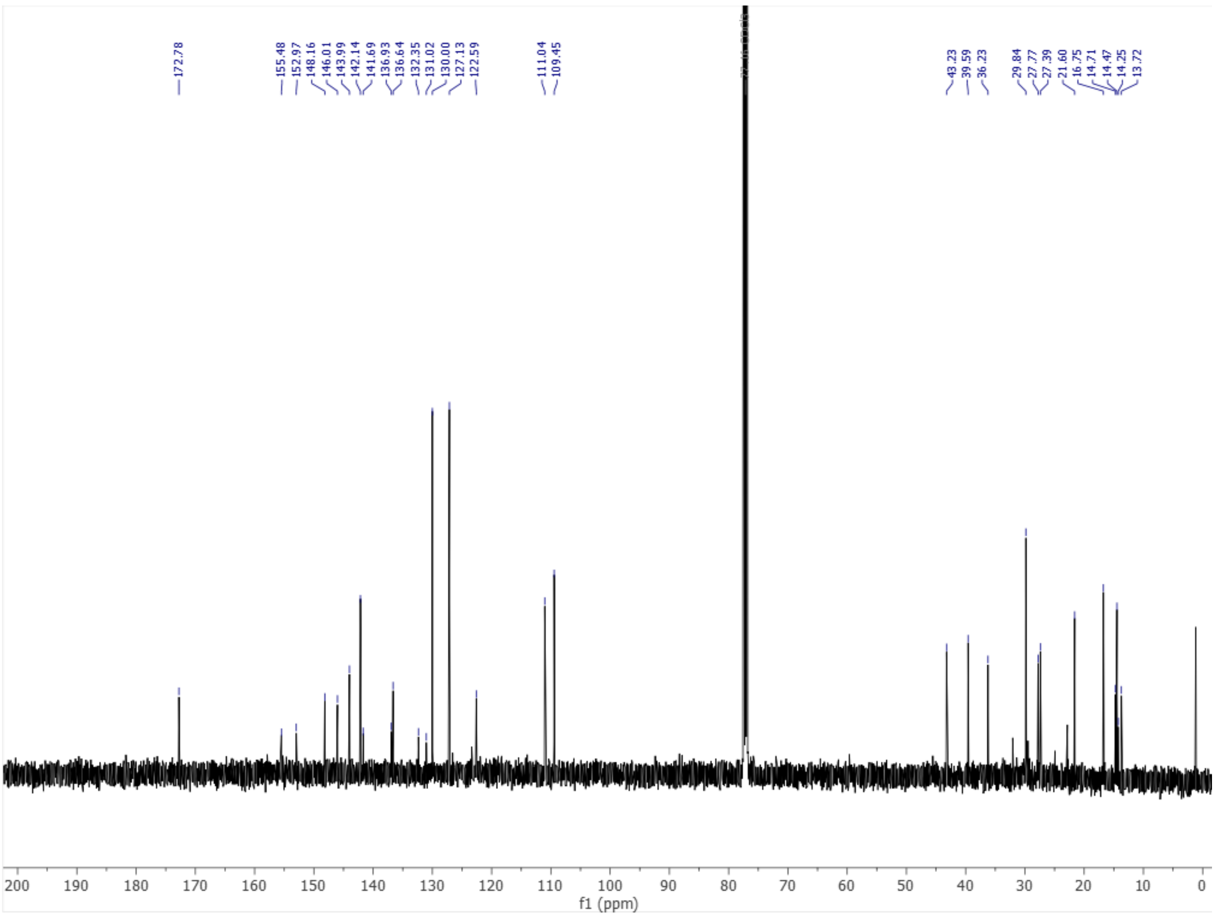


[13]C NMR spectrum of **PFB-ER**

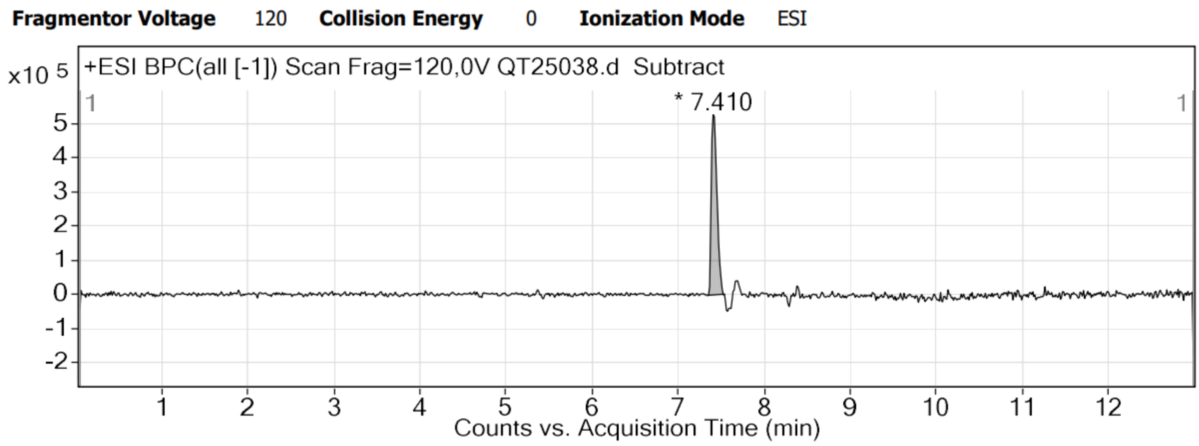


HPLC trace of **PFB-ER**

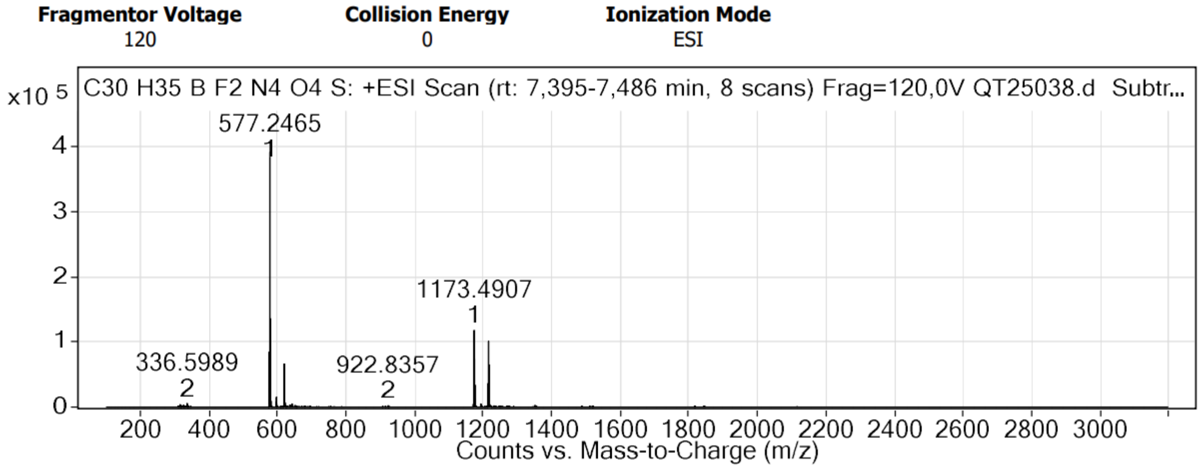


HRMS spectrum of **PFB-ER**

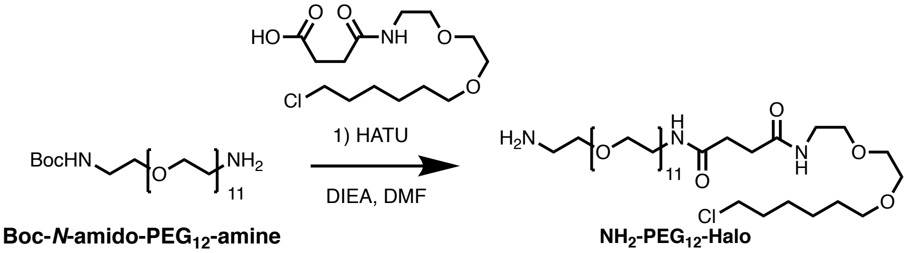


**$NH_2$-$PEG_{12}$-Halo**. To a solution of Boc-*N*-amido-$PEG_{12}$-amine (100 mg, 154 µmol, 1 eq) in DMF (2 mL) was added 4-[[2-[2-[(6-Chlorohexyl)oxy]ethoxy]ethyl]amino]-4-oxo-butanoic Acid (50 mg, 154 µmol, 1 eq), HATU (76 mg, 201 µmol, 1.3 eq) and DIEA (81 µL, 463 µmol, 3 eq). The solution was allowed to stir for 3 h at RT. The crude was concentrated under reduced pressure and purified by column chromatography on silica gel (DCM/MeOH: 9/1 to 8/2). Surprisingly, the **$NH_2$-$PEG_{12}$-Halo** was obtained (50 mg, 38%) as free amine, where the Boc protecting has been removed. $^1$H NMR (400 MHz, $CDCl_3$) δ 7.90 (s, 2H), 6.90 (t, J = 5.6 Hz, 1H), 6.64 (t, J = 5.6 Hz, 1H), 3.82 – 3.75 (m, 2H), 3.67 (dd, J = 5.6, 2.7 Hz, 2H), 3.65 – 3.46 (m, 50H), 3.46 – 3.33 (m, 6H), 3.15 (t, J = 4.9 Hz, 2H), 2.48 (s, 4H), 1.80 – 1.68 (m, 2H), 1.57 (p, J = 6.8 Hz, 2H), 1.48 – 1.27 (m, 4H). $^{13}$C NMR (101 MHz, $CDCl_3$) δ 71.23, 70.37, 70.35, 70.29, 70.26, 70.21, 70.19, 70.11, 70.05, 70.01, 69.99, 69.96, 69.87, 69.82, 69.69, 67.07, 45.03, 39.98, 39.21, 32.50, 31.68, 31.55, 29.42, 26.65, 25.38. HRMS (ESI+) calculated for $C_{38}H_{77}ClN_3O_{15}$ $[M+H]^+$ 850.5043, found 850.5036.

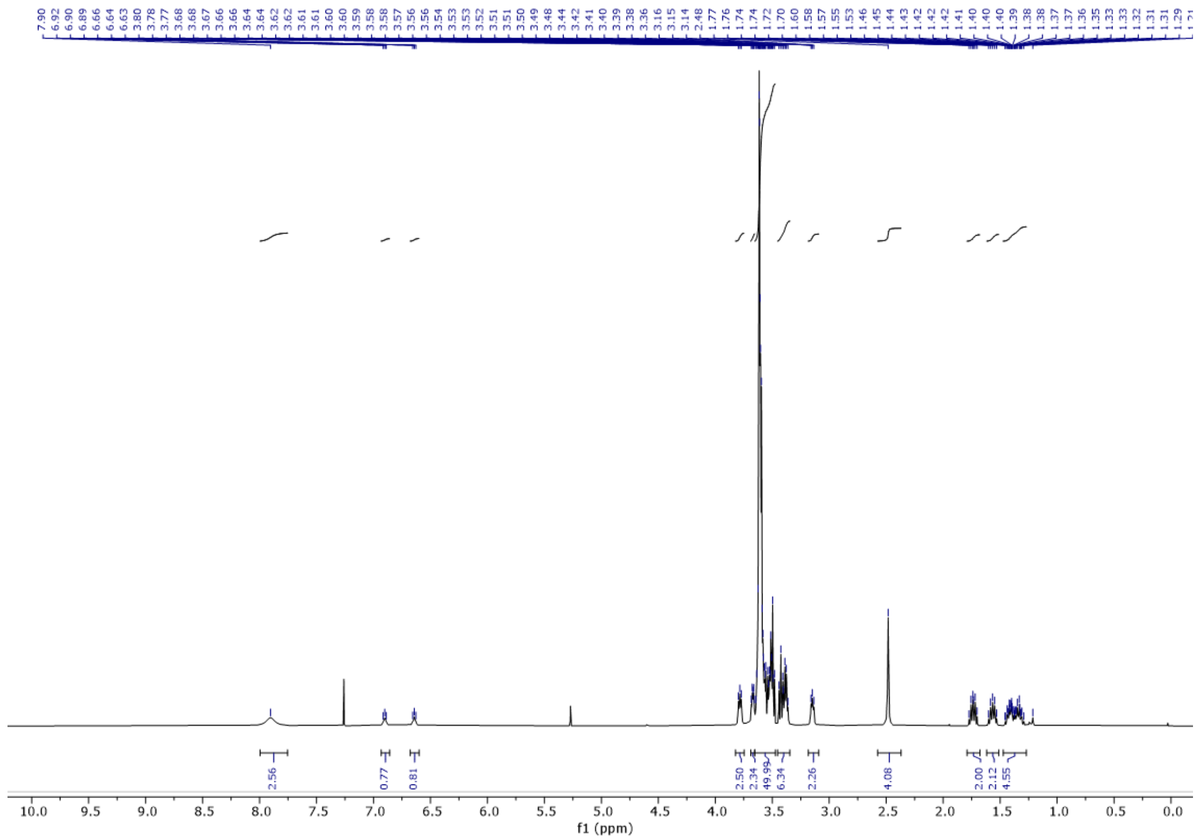

$^{1}H$ NMR spectrum of **$NH_2$-PEG$_{12}$-Halo**

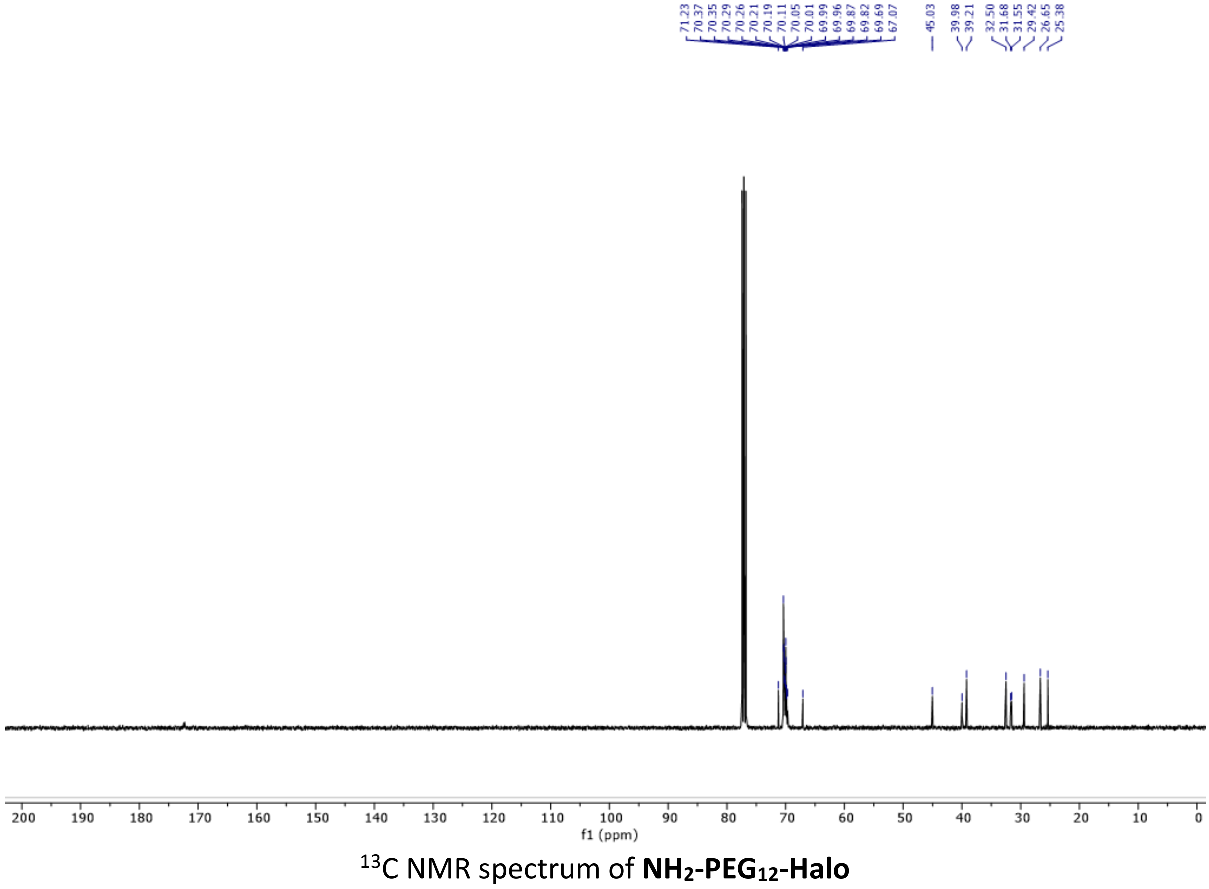

$^{13}C$ NMR spectrum of **$NH_2$-PEG$_{12}$-Halo**

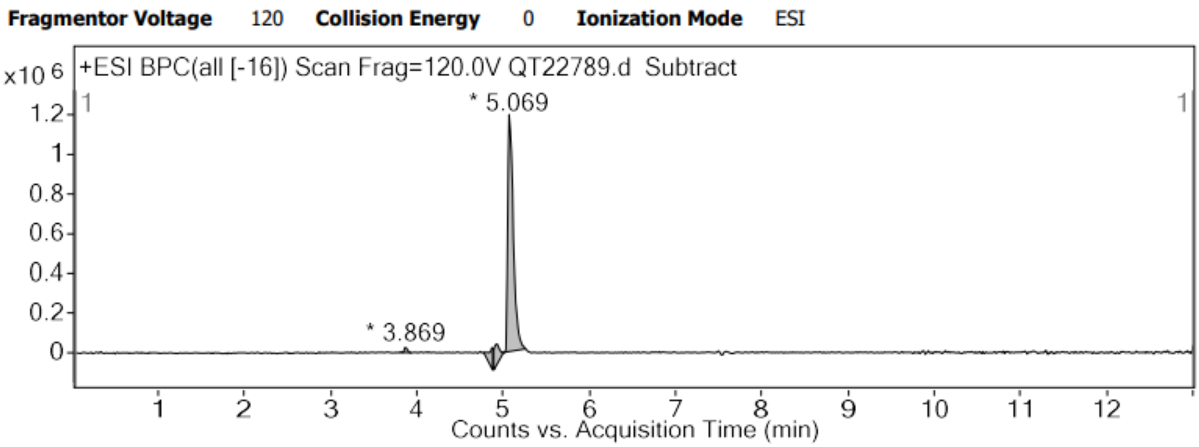


HPLC trace of **$NH_2$-$PEG_{12}$-Halo**

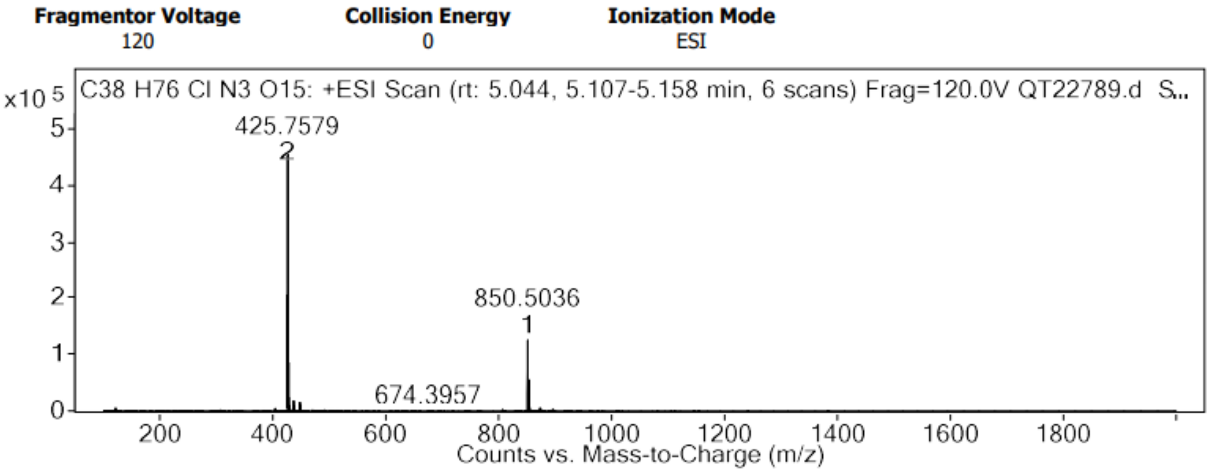


HRMS spectrum of **$NH_2$-$PEG_{12}$-Halo**

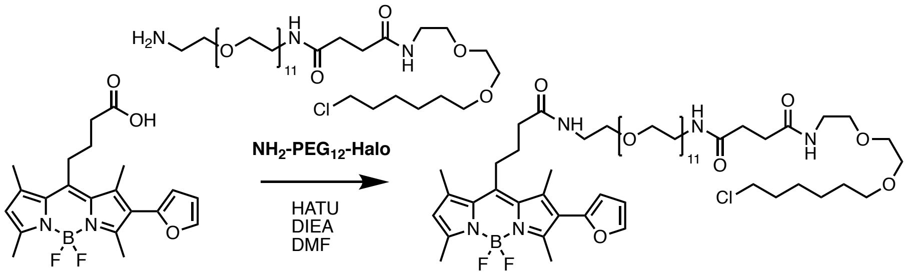


4 PFB-Halo

**PFB-Halo**. To a solution of **4** (5 mg, 12.5 µmol, 1 eq) in DMF (3 mL) was added **$NH_2$-$PEG_{12}$-Halo** (10.6 mg, 12.5 µmol, 1 eq), HATU (6.4 mg, 16.2 µmol, 1.3 eq) and DIEA (6.5 µL, 37 µmol, 3 eq). The solution was left to stir for 3 hours. The crude was concentrated under reduced pressure and purified by column chromatography on silica gel (DCM/MeOH: 9/1) to obtain **PFB-Halo** (9 mg, 58%). Rf = 0.50 (DCM/MeOH : 9/1). $^1$H NMR (400 MHz, $CDCl_3$) δ 7.49 (s, 1H, Furan), 6.65 (s, 1H, NH), 6.52 (s, 1H, NH), 6.48 (s, 1H, Furan), 6.30 (d, *J* = 3.2 Hz, 1H, Furan), 6.08 (s, 1H, ArH BODIPY), 3.84 – 3.32 (m, 56H), 3.15 – 3.01 (m, 2H), 2.61 – 2.42 (m, 16H, Methyl BODIPY and Alkyl PEG-Halo), 2.37 (t, *J* = 7.0 Hz, 2H), 2.02 – 1.95 (m, 2H), 1.76 (p, *J* = 7.0 Hz, 2H), 1.59 (p, *J* = 6.9 Hz, 2H), 1.48 – 1.33 (m, 4H). HRMS (ESI+) calculated for $C_{67}H_{103}BClF_2N_7NaO_{18}$ $[M+Na]^+$ 1254.6527, found 1254.6225

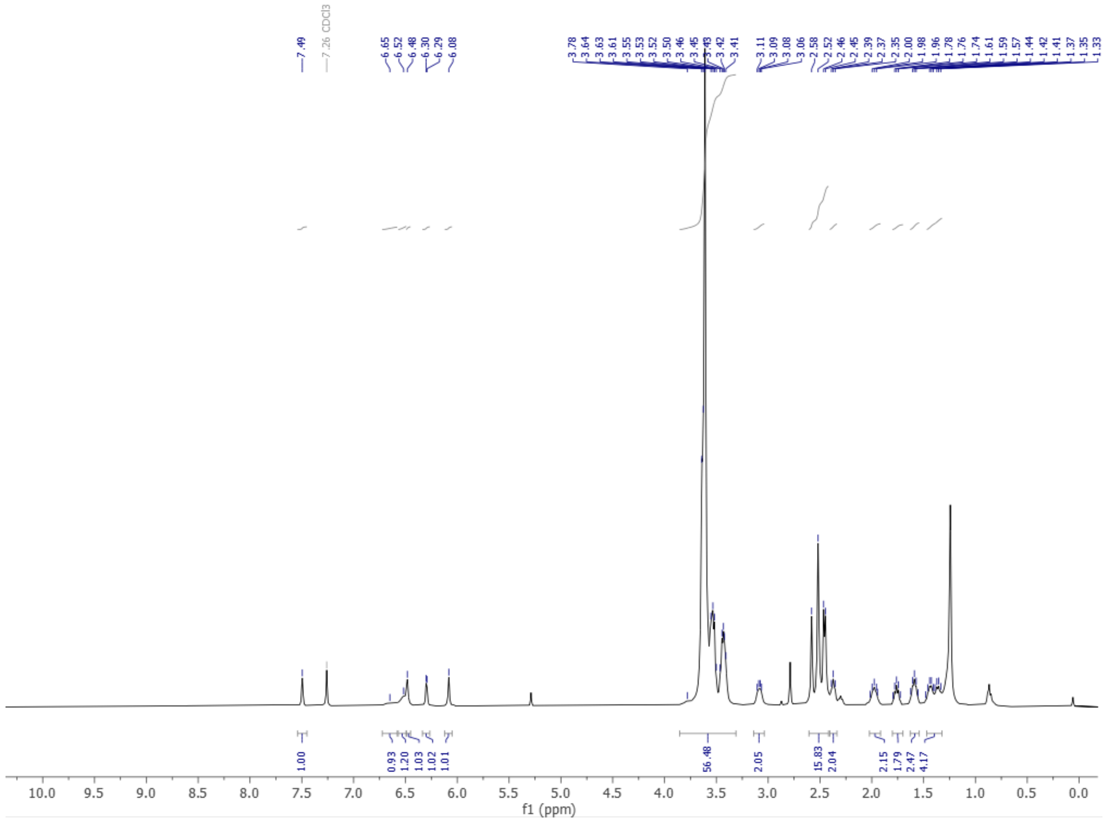


$^{1}$H NMR spectrum of **PFB-Halo**

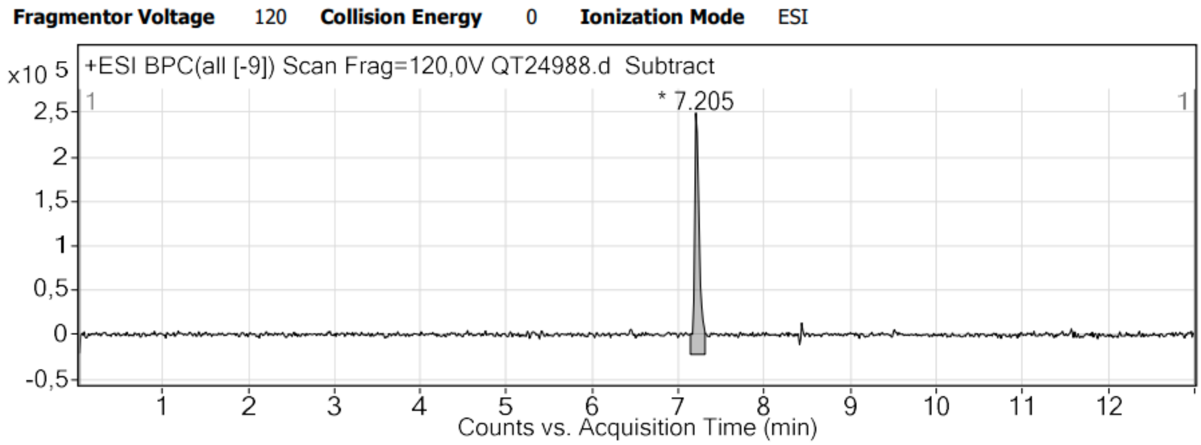


HPLC trace of **PFB-Halo**

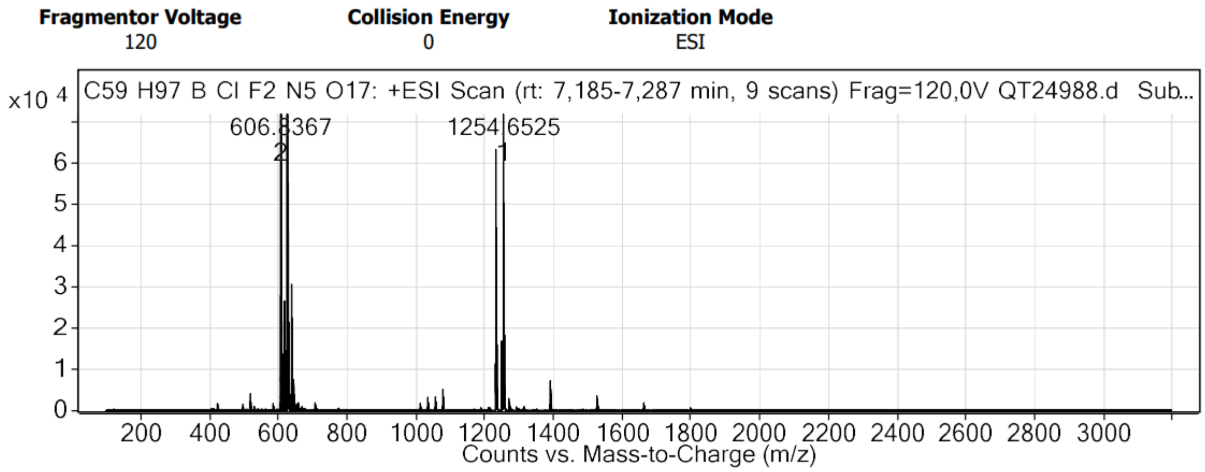


HRMS spectrum of **PFB-Halo**

**- Spectroscopy.** The water used for spectroscopy was Milli-Q water (Millipore), and all the solvents were spectroscopy grade. Absorption and emission spectra were recorded on a Cary 4000-HP spectrophotometer (Varian) and a FluoroMax-4 spectrofluorometer (Horiba Jobin Yvon) equipped with a thermo-stated cell compartment, respectively. For standard recording of fluorescence spectra, the emission was collected 10 nm after the excitation wavelength. All the spectra were corrected from the wavelength-dependent response of the detector. The fluorescence quantum yields $\varphi_F$ were determined following the following equation:

$$\varphi_F = \varphi_{ref} \times \frac{\int I_{fluo}^{sample} \, d\lambda}{\int I_{fluo}^{ref} d\lambda} \times \frac{OD_{ref}}{OD_{sample}} \times \frac{n_{sample}^2}{n_{ref}^2}$$

With OD the optical density at the excitation wavelength, I the intensity of fluorescence, dλ the wavelength interval and $n$ refraction index of the solvent.

**Fluorescence Lifetime Measurements.** Time-resolved fluorescence measurements were performed using time-correlated single-photon counting technique. Excitation pulses at 510 nm for **PFB-Mito** and 500 nm for **MB-Mito** were generated by a supercontinuum laser (NKT Photonics SuperK Extreme) with 10 MHz repetition rate. The fluorescence decays were collected at 640 nm for **PFB-Mito** in, and 520 nm for **MB-Mito** and aPFB-Mito, using a polarizer set at magic angle and a 16 nm band-pass monochromator (Jobin Yvon). The single-photon events were detected with a microchannel plate photomultiplier R3809U Hamamatsu, coupled to a pulse pre-amplifier HFAC (Becker-Hickl GmbH) and recorded on a time-correlated single photon counting board SPC-130 (Becker-Hickl GmbH).
Time-resolved exponential decays were fitted by using the global fit procedure of Igor Pro (Wavemetrics). The fitting function was a sum of exponential decays (up to 4 components) convolved with a normalized Gaussian curve of standard deviation σ standing for the temporal IRF and a Heavy side function. All emission decays were fitted using a weighting that corresponds to the standard deviation of the photon number squared root.

**Photoactivation studies. Determination of photoactivation constants.** Laser spectroscopy and conversion were performed using 3 × 3 mm optical path length quartz cuvettes of 45 µL at 1 µM in methanol. Excitation was provided by a continuous wave laser diode (488 nm,

Oxxius, Lannion, France) at 160 mW.cm$^{-2}$ for activation and 140 mW.cm$^{-2}$ for bleaching experiments. Photons were detected by a QE pro spectrometer from Ocean Optics. All measurements were performed at room temperature. The kinetic rate of phototransformation, $k_{Pt}$, was determined by fitting the emission increase (integrated spectra) of the photoactivatable dye over time as described in Moerner's method[25] and according to the equation (1).

$$\text{Equation (1) } A(t) = A(1 - e^{(-k_{Pt}t)})$$

Where A(t) is the emission signal over time of the photoactivatable dye.

Then the quantum yield of phototransformation $\phi_{Pt}$ is given by the equation (2).

$$\text{Equation (2) : } \phi_{Pt} = \frac{k_{Pt}}{\sigma \cdot \frac{P\lambda}{Shc}}$$

Where $P$ is the power of the laser used for irradiation (W), λ the wavelength of irradiation (m), $S$ the irradiated surface (here S = 0.15 cm²), $h$ the Planck's constant, $c$ the celerity of light (m.s$^{-1}$) and $\sigma$ the absorption cross section (cm²) defined from the molar-absorption coefficient of the dye at the excitation wavelength $\varepsilon$ (L.mol$^{-1}$.cm$^{-1}$) and the Avogadro's constant $N_{AV}$ (mol$^{-1}$) by equation (3):

$$\text{Equation (3) : } \sigma = \frac{\varepsilon \cdot 2303}{N_{AV}}$$

Based on our model, we hypothesized that a**PFB-Mito** was similar to **MB-Mito**. The chemical yield (η) is obtained by comparing the fluorescence intensity of **PFB-Mito** after activation ($Fl_A$) and the fluorescence intensity of **MB-Mito** in the same conditions by the equation (4).

$$\text{Equation (4) : } \eta = \frac{Fl_B}{Fl_A}$$

Finally, the quantum yield of photoactivation ($\varphi_{Pc}$) was obtained by multiplying the quantum yield of phototransformation ($\varphi_{Pt}$) by the chemical yield following the equation (5) and finally the quantum yield of photobleaching ($\varphi_{Bl}$) is determined knowing $\varphi_{Pc}$ and $\varphi_{Pt}$ by the equation (6).

$$\text{Equation (5) : } \phi_{act} = \phi_{Pt}\,\eta$$
$$\text{Equation (6) : } \phi_{Pt} = \phi_{Bl} + \phi_{Act}$$

Consequently, when the dye is not activable the quantum yields of phototransformation and photobleaching are linked following the equation (7).

$$\text{Equation (7) : } \phi_{Pt} = \phi_{Bl}$$

The photobleaching of the activated form (**aPFB-Mito**) was performed at 488 nm (140 mW.cm$^{-2}$) (after the photoactivation step) and the quantum yield of photobleaching was determined as described in our previous work.[3]

**Singlet oxygen quantum yield**. A solution of DPBF (100 µM) and the fluorophore or reference (5 µM) in MeOH were irradiated over few seconds with continuous wave laser. The emission spectra are acquired to obtain the slope of the decrease of DPBF fluorescence intensity. The singlet oxygen quantum yield is given by the following equation:

$$\varphi_{\Delta Fluorophore} = \varphi_{\Delta Reference} \frac{p_{Fluorophore} F_{Reference}}{p_{Reference} F_{Fluorophore}}$$

With $p_x$ the decrease slope avec the compound x and Fx the corrected absorption given by the following equation:

$$F_x = 1 - 10^{-OD}$$

**Cellular culture**. Hela cells were incubated in Dulbecco's Modified Eagle Medium (1 g·L$^{-1}$ glucose) supplemented with 10% fetal bovine solution, 1% L-glutamine, and 1% antibiotic solution (penicillin−streptomycin) at 37 °C in a humidified atmosphere containing 5% $CO_2$. Cells were seeded onto a chambered coverglass (IBiDi) at a density of 1×10$^5$ cells/well 24 h before the microscopy measurement. For imaging, the culture medium was removed and the attached cells were washed with Opti-MEM (Gibco−Invitrogen).

**Confocal imaging.** Cells were incubated with **PFB-Mito** or **PFB-ER** (200 nM) in opti-MEM for 30 minutes. The cells were then washed with opti-MEM before being imaged. Cells were imaged with a Leica TSC SP8 laser scanning confocal microscope with a 63× objective.
mCherry and ER Tracker™ Red were exited at 560 nm and the fluorescence signal was collected from 570 to 700 nm. Membright®-640, Mitotracker™ DeepRed, were exited at 640 nm and the fluorescence signal was collected from 650 to 750 nm.

**Transfection**. Hela cells have been transfected 24 h after being seeded in Ibidi chambers. Transfection mix have been prepared by preparing extemporaneously solution A containing jetPEI® (2 µL, Polyplus transfection®) in 150 mM NaCl (50 µL) and solution B of Halo tagged protein-coding pDNA (1.6 µg) and pcDNA3.1-mCherry (0.2 µg) in 150 mM NaCl (50 µL). After 5 min at room temperature, solutions A and B were mixed thoroughly and stayed 5 min at room temperature. Transfection mix was added on cells with 1 mL growing media. Cells were incubated at 37 °C for 24 h. The medium was removed and the cells were washed twice with opti-MEM, then **PFB-Halo** (50 nm) was added in opti-MEM and incubated for 30 minutes. To remove non-specific labelling the medium was removed and washed twice with growing medium and left at 37°C for 1 hour. Then the medium was removed, washed twice with opti-MEM before imaging in opti-MEM.

**Colocalisation studies, Determination of Pearson's constants**

The colocalization studies were performed on live HeLa cells stained with PFB-Mito and PFB-ER and co-stained respectively with Mito-tracker™ DeepRed (50 nm) and ER-Tracker™ Red (200 nm) (thermofisher scientific). The Pearson's coefficients were obtained using the imageJ plugin: JaCoP.[4]

**Sequential Photoactivation in cells.** Cells were found using a co-labelling, MemBright 640 (200 nm) for PFB-Mito, Hoechst 33258 (5 µg.mL$^{-1}$) for PFB-ER. Due to heterogeneous labelling when using PFB-Halo and transfection, the cells were co-transfected with an 8-fold equivalent of DNA plasmid coding for cytosolic mCherry protein. This protocol helped to find Halo-transfected cells by screening mCherry positive cells in the red channel. A region of interest (ROI) was made on one cell and about 20 scans with 488 nm laser was applied. The activation process was monitored through the evolution of the intensity histogram, the irradiation scans were stopped once the histogram presents no more evolution. After single cell activation a zoom out was performed and an image was taken of the whole field of view.

**Live SMLM.** Super-resolution localization microscopy imaging was performed on a home-built setup based on a Nikon Eclipse Ti microscope with 100X 1.49 NA oil-immersion objective. The excitation was provided with a 488 nm laser (3.3 $W.cm^{-2}$). An acousto-optic tunable filter (AOTF; Opto-Electronic) was used to modulate the laser power. The signal was detected with on an EM-CCD camera from Hamamatsu (ImagEM). This laser power was sufficient to trigger the photoactivation of the probe, and thus to find the Halo-tagged protein expressing cells. Single cells were imaged by 10 successive acquisitions (every 30 s) of 100 frames with 13.9 ms integration time (EM-Gain 600). The reconstruction of the 10 acquisitions provided dynamic super-resolution movie.

**SMLM treatments (Thunderstorm, SRRF plugin)**

**SMLM.** All images were analyzed using ThunderSTORM plugin.[5] Images were analyzed with a Wavelet filter (B-Spline) (order 3, scale 1). Localization of molecules was performed with a local maximum method using standard deviation, connectivity, 8-neighbourhood. Sub-pixel localization was done using a PSF of integrated Gaussian with fitting radius of 3 pixels and initial sigma of 1.6 pixels. The localizations were filtered with density filter (3 locks in 50 nm diameter) and by removing localizations with uncertainty of localization superior to 100 nm.

**SRRF.** The same images were also analyzed with SRRF plugin[6] (Ring radius : 0.5, Radiality Magnification : 6, Axes in Ring : 6).

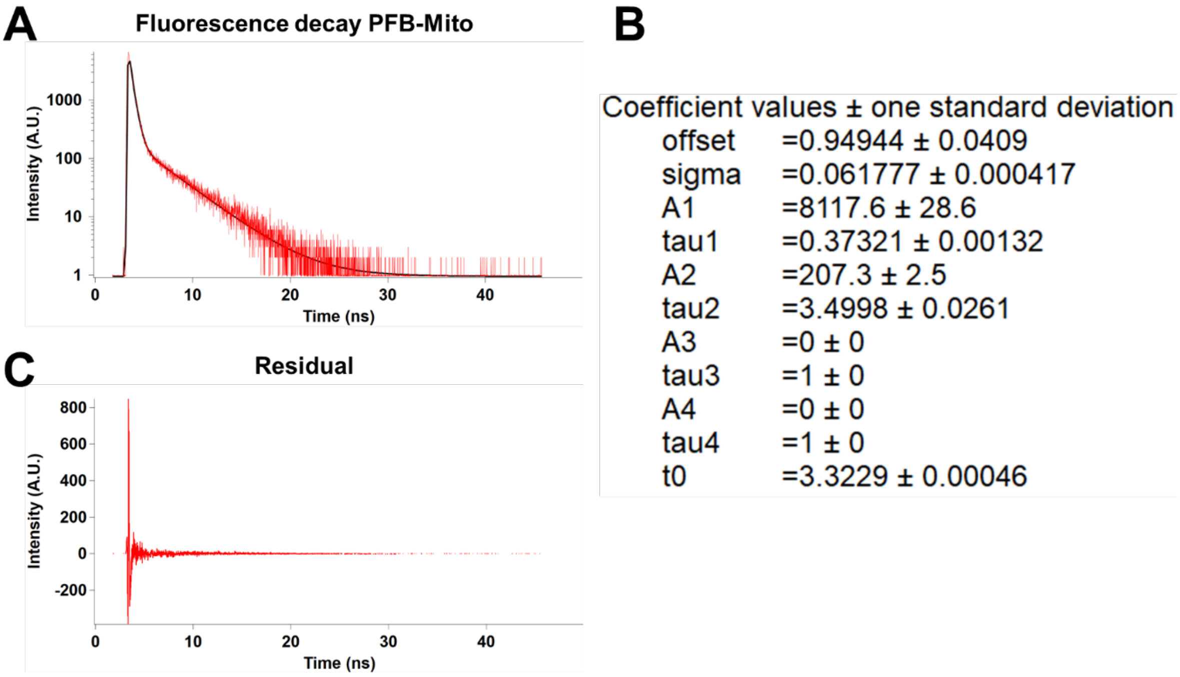


**Figure S1. Fluorescence decay of PFB-Mito**. (A) fluorescence decay of PFB-Mito, excitation at 510 nm and emission at 640 nm. (B) constant collected from the fit, the second exponential correspond to the small number of activated molecules. (C) residuals from the fit (not normalized).

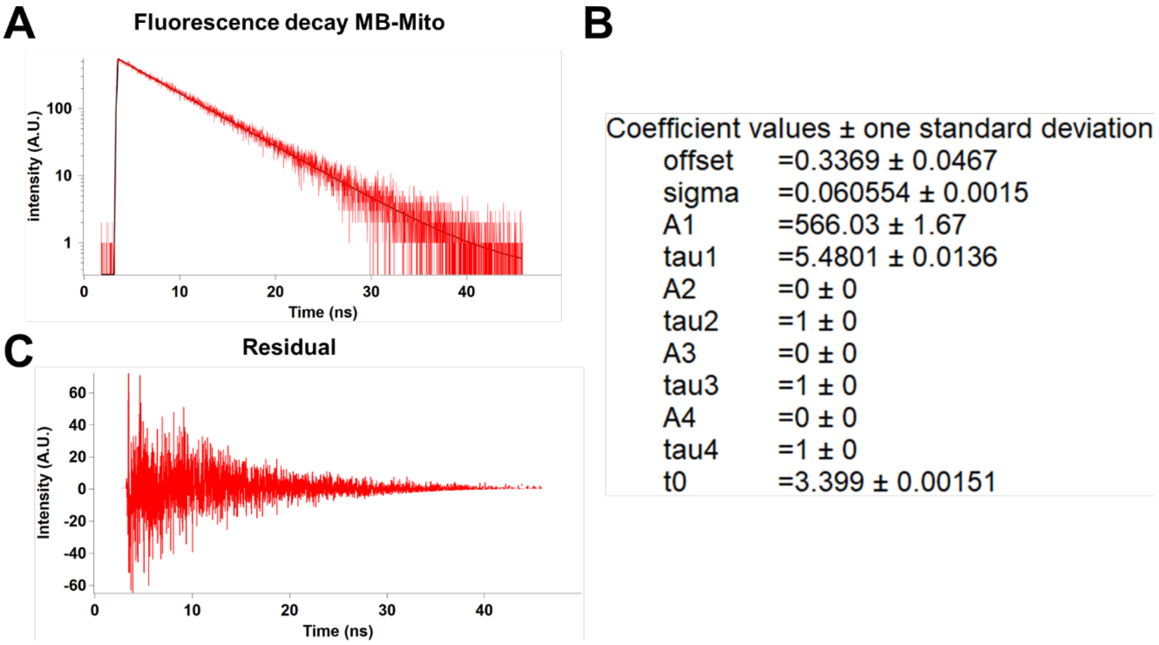


**Figure S2. Fluorescence decay of MB-Mito**. (A) fluorescence decay of MB-Mito, excitation at 500 nm and emission at 520 nm. (B) constant collected from the fit. (C) residuals from the fit (not normalized).

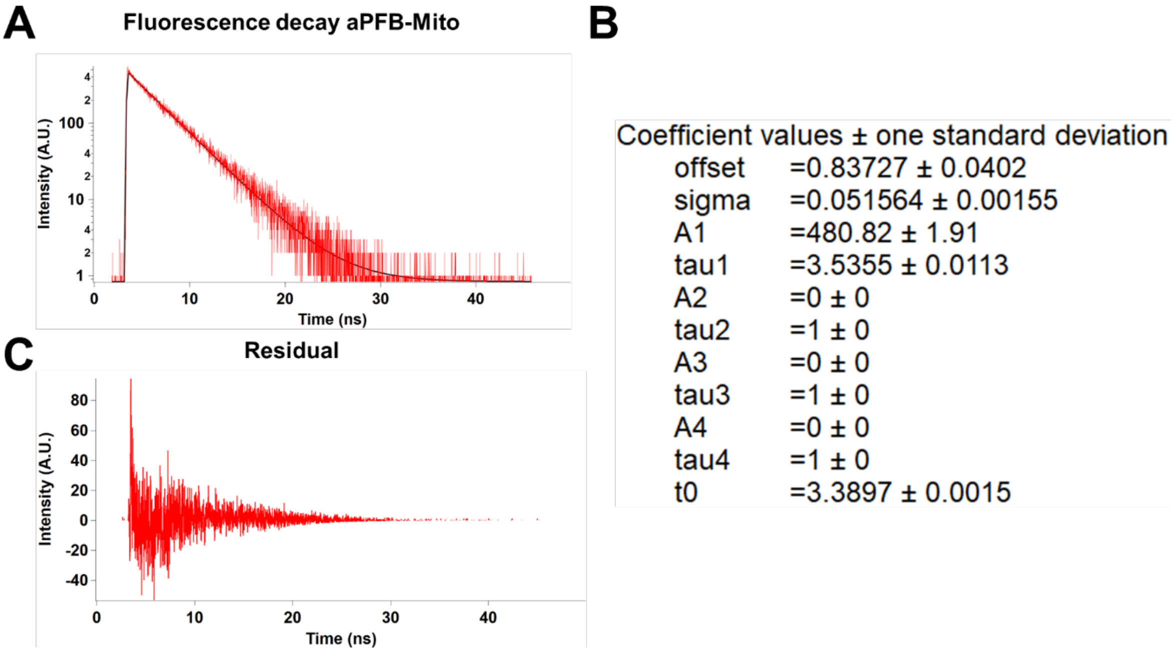


**Figure S3. Fluorescence decay of aPFB-Mito**. (A) fluorescence decay of aPFB-Mito, excitation at 500 nm and emission at 520 nm. (B) constant collected from the fit. (C) residuals from the fit (not normalized).

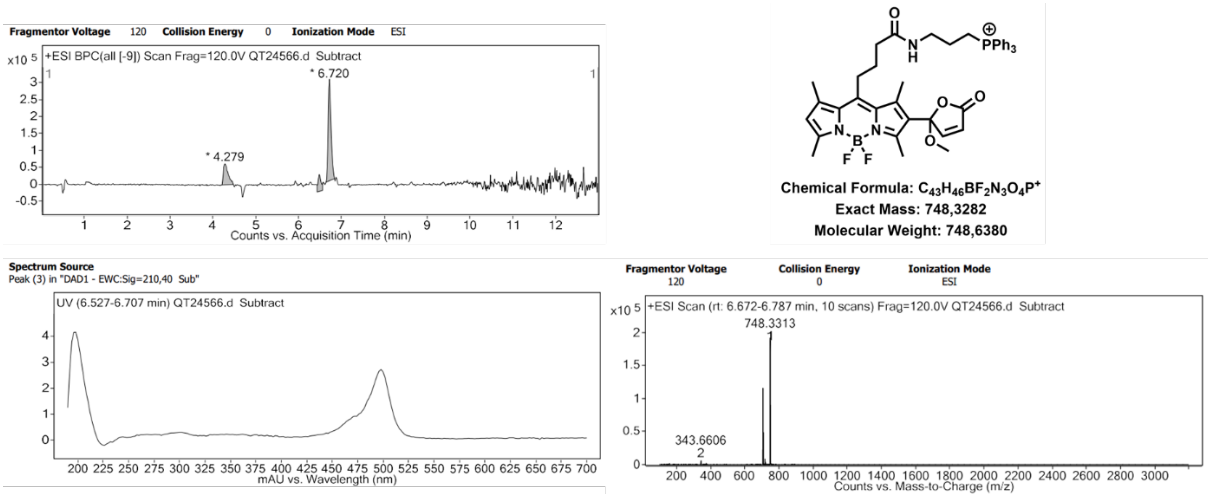


**Figure S4. HRMS analysis of the major photoproduct cPFB-Mito.** The results showed a mass as well as an absorbance spectrum in accordance to the expected photoproduct **cPFB-Mito**.

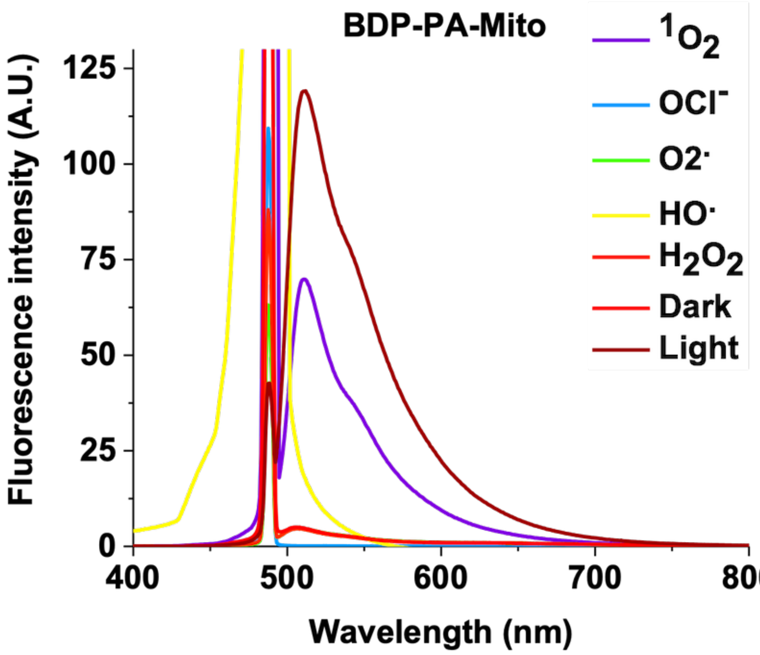


**Figure S5.** Emission spectra of **PFB-Mito** in the presence of various ROS and without ROS in the dark (negative control). Selectivity tests were performed adding various ROS generators in water in a methanolic solution of **PFB-Mito** (1 µM). $^1O_2$: 5 µM of Aluminium Phthalocyanine with 638 nm laser for 30 minutes: $O_2^{\bullet -}$ : 2 mM of $KO_2$ for 30 minutes in the dark; OCl- : 2 mM of NaOCl for 30 minutes in the dark; $^{\bullet}OH$: 2 mM of $FeSO_4$ + 2 mM $H_2O_2$ for 30 minutes in the dark; $H_2O_2$: 2 mM of $H_2O_2$ for 30 minutes in the dark. The emission spectra were acquired upon 488 nm laser excitation (4 $mW.cm^{-2}$).

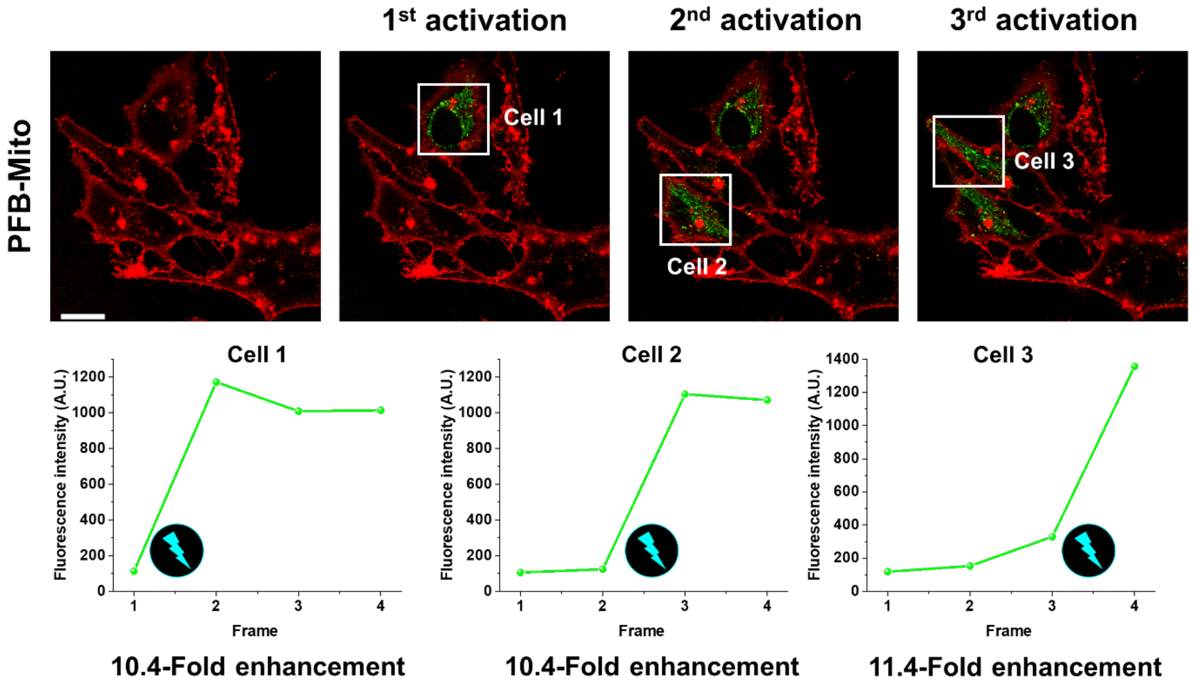


**Figure S6. Sequential activation of live cells mitochondria using PFB-Mito**. (Top) Laser scanning confocal images of Hela cells stained with PFB-Mito (200 nM). The cells were sequential irradiated using the 488 nm laser line to activate the PFB-Mito-stained mitochondria (green). The plasma membrane (red) was stained with MemBright-640 (200 nm). Scale bar is 20 µm. (Bottom) mean intensity of the cells before and after each activation. The interval between activation was 20 s.

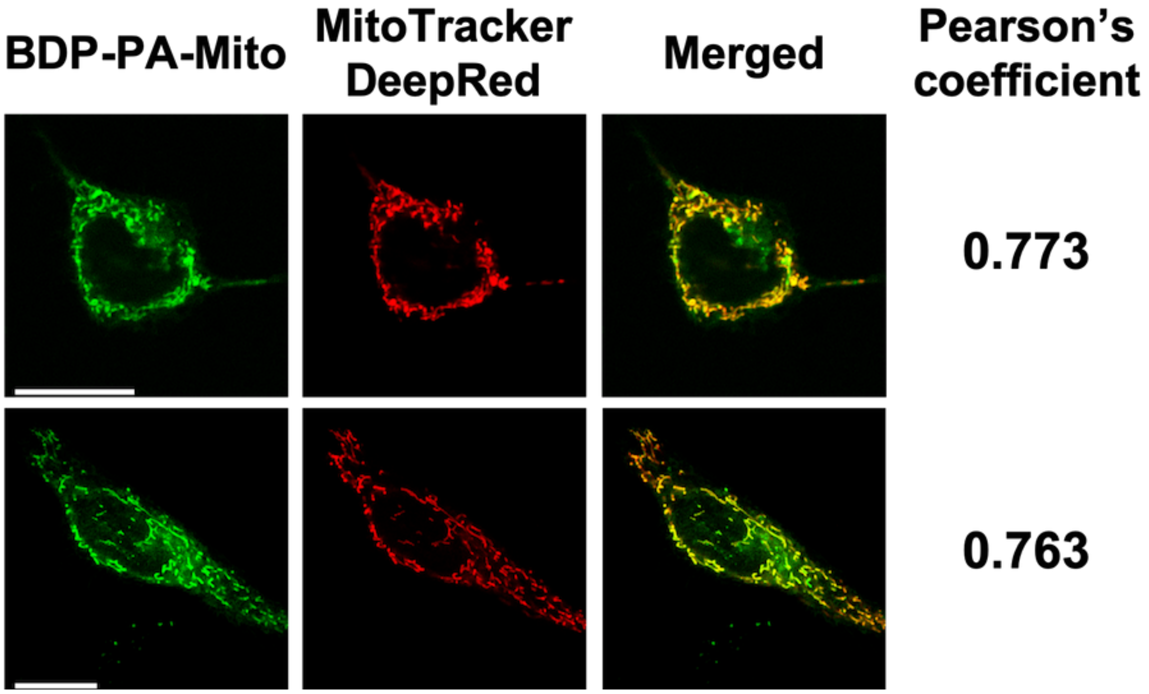


**Figure S7.** Laser scanning confocal images of HeLa cells stained with PFB-Mito (200 nM) and Mitotracker deep-red (50 nM). Prior to colocalization studies, the cells were exposed to the laser line at 488 nm to trigger the photoactivation of PFB-Mito. Scale bar is 20 µm.

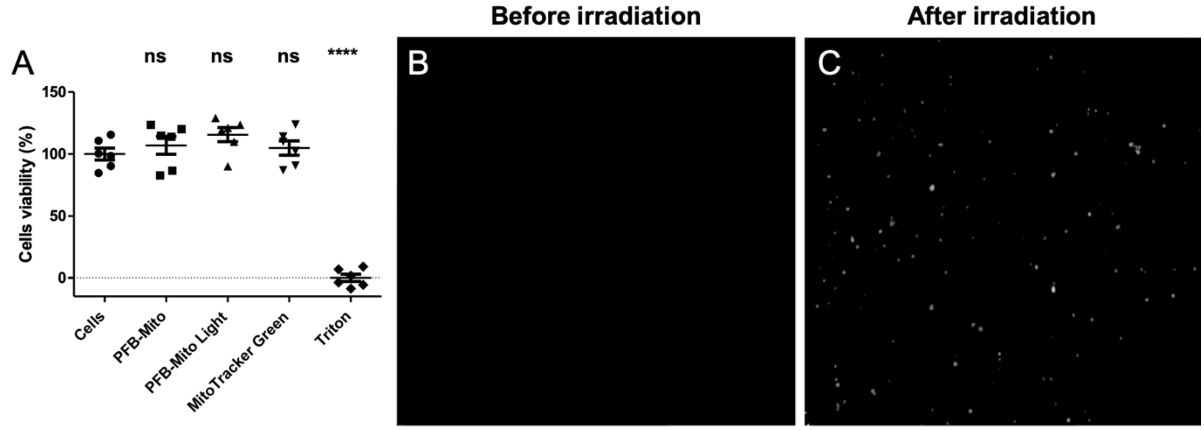


**Figure S8.** Cyto-and photocytotoxicity of PFB-Mito. (A) MTT assays of Hela cells incubated with PFB-Mito or Mito-Tracker Green (200 nM), for 30 min. The results showed no significant cytotoxicity (PFB-Mito) nor photocytotoxicity (PFB-Mito Light) of both probes. (B)Widefield microscopy image of cells incubated with PFB-Mito before activation and before the MTT test. (C) Brightfield microscopy image of cells incubated with PFB-Mito after exposure to light and before the MTT test proving that the activation occurred.

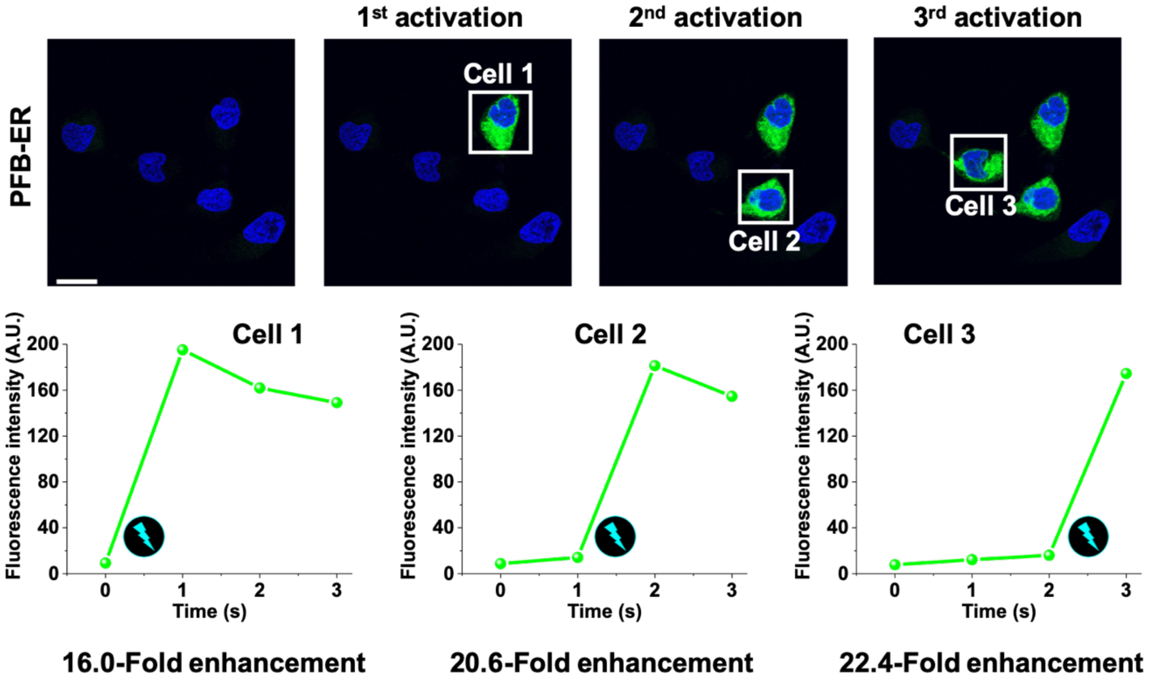


**Figure S9. Sequential activation of live cells with PFB-ER.** (Top) Laser scanning confocal images of Hela cells' endoplasmic reticulum stained with PFB-ER (200 nM). The cells were sequentially irradiated using the 488 nm laser line to activate the PFB-ER-stained endoplasmic reticulum (green). The nucleus (blue color) was stained with Hoechst 33258 (5 µg. mL$^{-1}$). Scale bar is 20 µm. (Bottom) mean intensity of the cells before and after each activation. The interval between activation was 20 s.

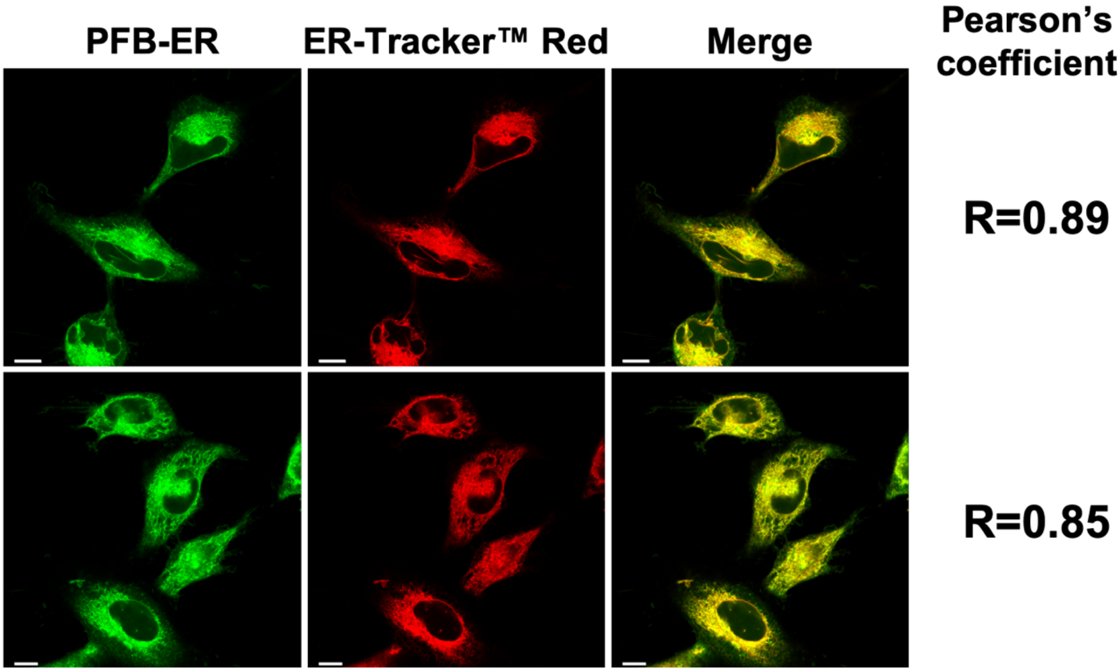


**Figure S10.** Laser scanning confocal images of HeLa cells stained with PFB-ER (200 nM) and ER-Tracker™ Red (200 nM). Prior to colocalization studies, the cells were exposed to the microscope's lamp at 488 nm to trigger the photoactivation of PFB-ER. Scale bar is 10 µm.

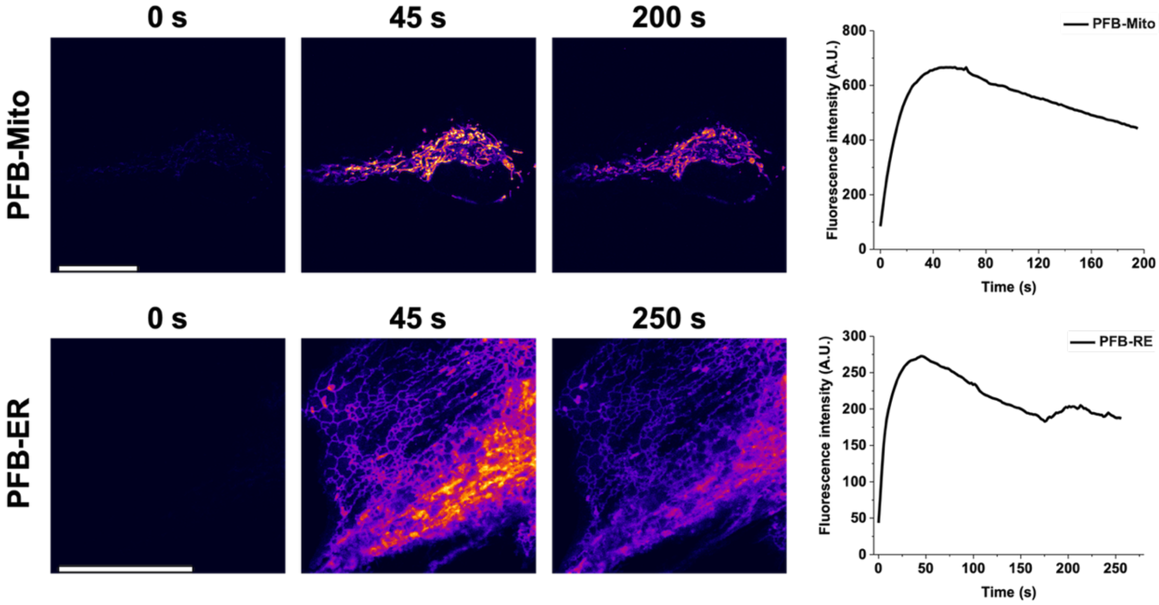


**Figure S11. Photoactivation and fatigue resistance of PFB-Mito and PFB-ER.** Laser scanning confocal images of Hela cells stained with PFB-Mito and PFB-ER (200 nM) over time (left). Fluorescence intensity individual cells showing the fast activation followed by the slow decay. (right). Individual cells were found and targeted for activation using plasma membrane staining with MemBright-640. Scale bar is 20 µm.

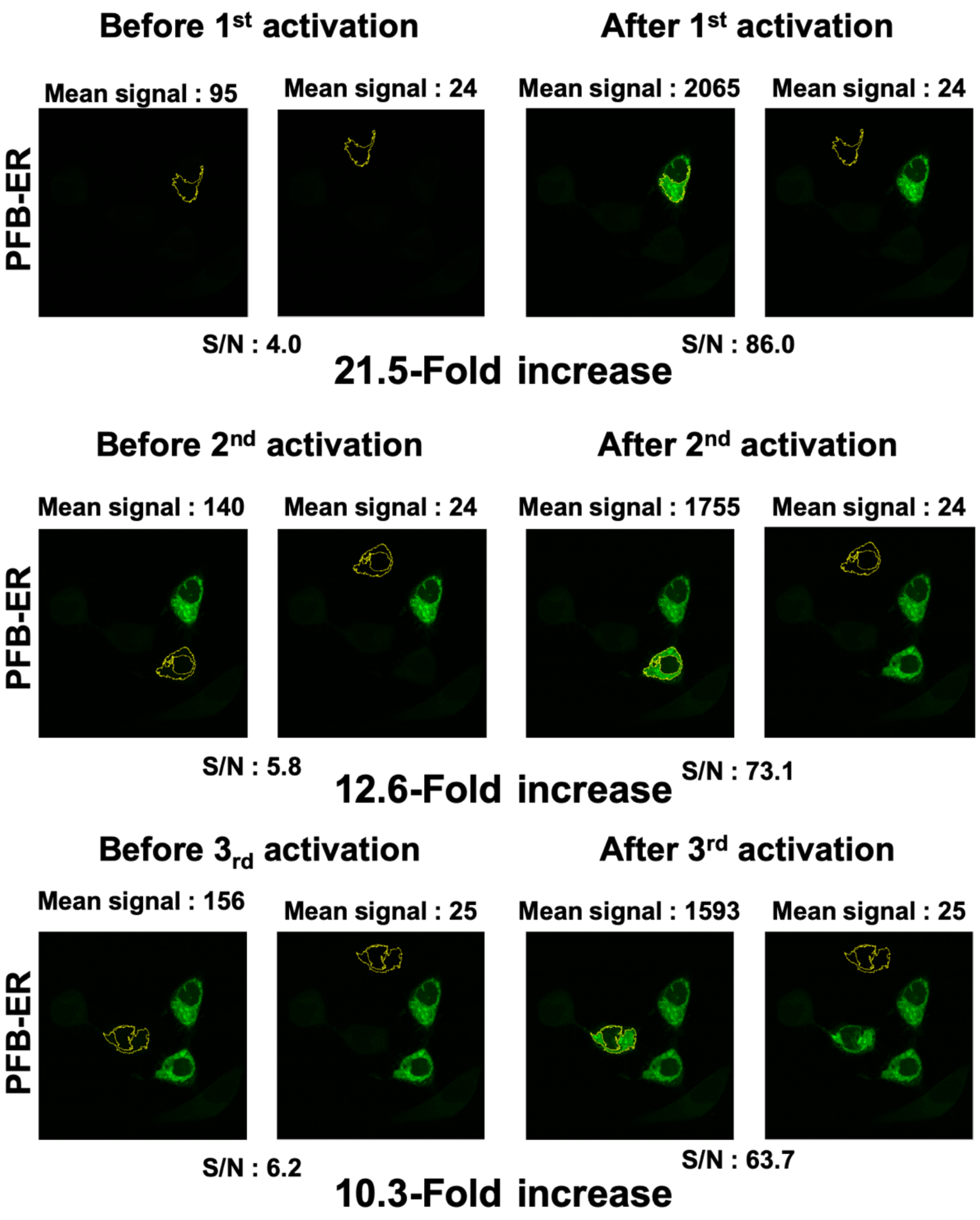


**Figure S12. Determination of the signal to noise ratio in imaging example of PFB-ER.** Once PFB activated, a region of interest (ROI) is made to monitor the signal (right). The same ROI is moved on a dark region to get the S/N signal. The same ROI was applied on the same cell before activation of PFB to obtain the S/N signal before activation (left). This analysis was performed on three different cells.

**Table S1.** Signal to noise ratio (S/N) of PFB in various sub-cellular environments before and after activation.

| Dye | S/N | S/N Activated | Enhancement fold |
|---|---|---|---|
| **PFB-Mito** | 2 ± 0 | 17 ± 2 | 9 ± 1 |
| **PFB-RE** | 5.3 ± 1.2 | 77.6 ± 12.1 | 14.7 ± 6.0 |
| **PFB-Halo Actin** | 6.8 ± 5.8 | 20.1 ± 13.2 | 3.3 ± 0.7 |
| **PFB-Halo H2B** | 6.7 ± 2.5 | 18.0 ± 10.8 | 2.5 ± 0.7 |
| **PFB-Halo Nucl** | 37.6 ± 2.5 | 87.4 ± 12.6 | 2.3 ± 0.4 |

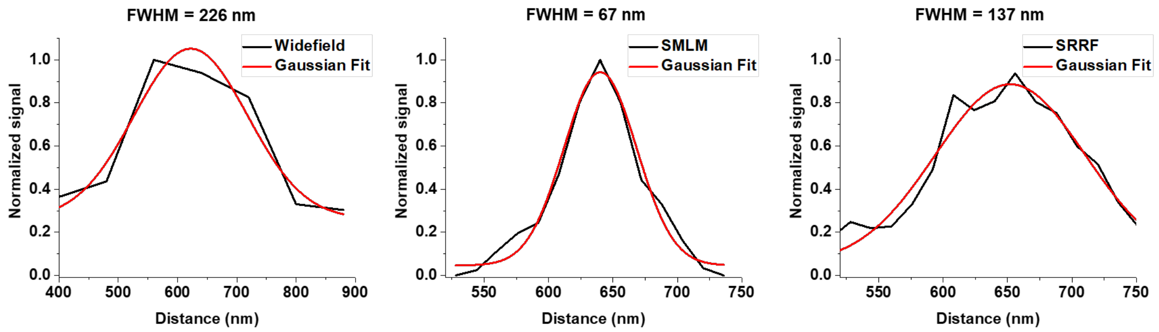


**Figure S13.** Plot profile corresponding to the ROI line trace in figure 5D for widefield, SMLM and SRRF images. A Gaussian fit was applied and the FWHM was determine to compare the obtained resolutions with the different imaging and treatment modes.

### References

(1) Janssen, A. P. A.; van der Vliet, D.; Bakker, A. T.; Jiang, M.; Grimm, S. H.; Campiani, G.; Butini, S.; van der Stelt, M. Development of a Multiplexed Activity-Based Protein Profiling Assay to Evaluate Activity of Endocannabinoid Hydrolase Inhibitors. *ACS Chem. Biol.* **2018**, *13* (9), 2406–2413. https://doi.org/10.1021/acschembio.8b00534.

(2) Thompson, M. A.; Biteen, J. S.; Lord, S. J.; Conley, N. R.; Moerner, W. E. Molecules and Methods for Super-Resolution Imaging. In *Methods in Enzymology*; Elsevier, 2010; Vol. 475, pp 27–59. https://doi.org/10.1016/S0076-6879(10)75002-3.

(3) Saladin, L.; Breton, V.; Dal Pra, O.; Klymchenko, A. S.; Danglot, L.; Didier, P.; Collot, M. Dual-Color Photoconvertible Fluorescent Probes Based on Directed Photooxidation Induced Conversion for Bioimaging. *Angewandte Chemie International Edition* **2023**, *62* (4), e202215085. https://doi.org/10.1002/anie.202215085.

(4) Bolte, S.; Cordelières, F. P. A Guided Tour into Subcellular Colocalization Analysis in Light Microscopy. *Journal of Microscopy* **2006**, *224* (3), 213–232. https://doi.org/10.1111/j.1365-2818.2006.01706.x.

(5) Ovesný, M.; Křížek, P.; Borkovec, J.; Švindrych, Z.; Hagen, G. M. ThunderSTORM: A Comprehensive ImageJ Plug-in for PALM and STORM Data Analysis and Super-Resolution Imaging. *Bioinformatics* **2014**, *30* (16), 2389–2390. https://doi.org/10.1093/bioinformatics/btu202.

(6) Culley, S.; Tosheva, K. L.; Matos Pereira, P.; Henriques, R. SRRF: Universal Live-Cell

Super-Resolution Microscopy. *The International Journal of Biochemistry & Cell Biology* **2018**, *101*, 74–79. https://doi.org/10.1016/j.biocel.2018.05.014.
